# A catalog of microbial genes from the bovine rumen unveils a specialized and diverse biomass-degrading environment

**DOI:** 10.1101/272690

**Authors:** Junhua Li, Huanzi Zhong, Yuliaxis Ramayo-Caldas, Nicolas Terrapon, Vincent Lombard, Gabrielle Potocki-Veronese, Jordi Estellé, Milka Popova, Ziyi Yang, Hui Zhang, Fang Li, Shanmei Tang, Weineng Chen, Bing Chen, Jiyang Li, Jing Guo, Cécile Martin, Emmanuelle Maguin, Xun Xu, Huanming Yang, Jian Wang, Lise Madsen, Karsten Kristiansen, Bernard Henrissat, Stanislav D. Ehrlich, Diego P. Morgavi

## Abstract

**Background:** The rumen microbiota provides essential services to its host and, through its role in ruminant production, contributes to human nutrition and food security. A thorough knowledge of the genetic potential of rumen microbes will provide opportunities for improving the sustainability of ruminant production systems. The availability of gene reference catalogs from gut microbiomes has advanced the understanding of the role of the microbiota in health and disease in humans and other mammals. In this work, we established a catalog of reference prokaryote genes from the bovine rumen.

**Results:** Using deep metagenome sequencing we identified 13,825,880 non-redundant prokaryote genes from the bovine rumen. Compared to human, pig and mouse gut metagenome catalogs, the rumen is larger and richer in functions and microbial species associated with the degradation of plant cell wall material and production of methane. Genes encoding enzymes catalyzing the breakdown of plant polysaccharides showed a particularly high richness that is otherwise impossible to infer from available genomes or shallow metagenomics sequencing. The catalog expands by several folds the dataset of carbohydrate-degrading enzymes described in the rumen. Using an independent dataset from a group of 77 cattle fed 4 common dietary regimes, we found that only <0.1% of genes were shared by all animals, which contrast with a large overlap for functions, i.e. 63% for KEGG functions. Different diets induced differences in the relative abundance rather than the presence or absence of genes explaining the great adaptability of cattle to rapidly adjust to dietary changes.

**Conclusions:** These data bring new insights into functions, carbohydrate-degrading enzymes and microbes of the rumen that is complementing the available information on microbial genomes. The catalog is a significant biological resource enabling deeper understanding of phenotypes and biological processes and will be expanded as new data is made available.

## Background

Ruminant production contributes to livelihood and to food and nutritional security in many regions of the world. Milk and meat from ruminants are important sources of protein and micronutrients in the human diet but often criticized as unsustainable because of the low conversion efficiency of plant feeds into animal foods [1] and also due to the high environmental footprint. However, when the feed conversion efficiency of protein and energy contained in milk and meat is calculated based on the ingestion of human-inedible protein and energy the output is higher that the input, particularly in forage-based production systems [2, 3]. The transformation of feeds, not suitable for human consumption, into highly nutritious protein and energy, products is carried out by gastrointestinal symbiotic microbes, particularly those residing in ruminants’ forestomach –the rumen. Rumen microbes are essential for ruminants allowing them to thrive in agricultural land not suitable for crops and to consume agricultural byproducts unfit for other livestock species. The enhanced functions provided by the rumen microbiota are key for the characteristic adaptability and robustness of ruminants to cope with nutritional and climatic stresses [4].

Improving our understanding of the rumen microbiota provides opportunities for knowledge-based strategies aiming at enhancing efficacy in ruminant production while minimizing its negative effect on the environment. Great advances on microbiota functions in the rumen has been obtained by extensive genome sequencing of cultured rumen bacteria and archaea (Hungate1000 project) [5] and by assembling of draft genomes from metagenomic data [6, 7]. These catalogs of reference genomes and metagenome-assembled genomes (MAGs) give great insight into the functionality of this ecosystem but, however extensive, they still do not cover the full bacterial and archaeal diversity present in the rumen [8–10]. In this study, we used a complementary approach to generate a catalog of unique rumen prokaryotic genes that enabled us to decipher functional potentials of the microbiota as a whole, in particular the capacity to deconstruct structural carbohydrates from forages, and we explored the effect of feed on the microbiota composition and functions.

## Data description

### Construction of a bovine rumen prokaryotic gene catalog

To build a bovine rumen prokaryotic gene catalog, we collected samples of total rumen content samples from five Holstein cows and five Charolais bulls. For reducing the ecosystem complexity and to improve metagenome assemblies, rumen ciliated protozoa were depleted from the samples before microbial DNA extraction. A total of 1,206 Gb of raw metagenomic sequencing data were generated with an average 111 Gb clean data for each animal. This sequencing depth, much greater than that used for gut gene catalogs from humans and other monogastric animals [11–13] was necessary to enable the assembly of the more complex rumen microbiome. After *de novo* assembly, open reading frames (ORF) prediction and removal of redundancy, 13,825,880 non-redundant genes were obtained with an average length of 716 base pairs (bp) and 39% of these genes were complete ORFs (Supplementary Table 1).

Compared to the large rumen gene catalog published by Hess et al. [14], the number of non-redundant genes discovered in this study is more than 5 fold larger; shared genes were in most cases also longer (Figure 1a & b and Supplementary Table 2). Thus, the mapping rate of reads from 77 additional rumen samples obtained in this study and eight published rumen samples from UK cattle [15] increased from ∼10% using the previous catalog [14] to ∼40% (11-51%) (Supplementary Figure 1 a & b). This confirms that the representativeness of the rumen catalog was greatly improved, even though the mapping efficiency was still relatively low, as compared to 80% for the human gut microbiome [11].

**Figure 1.**
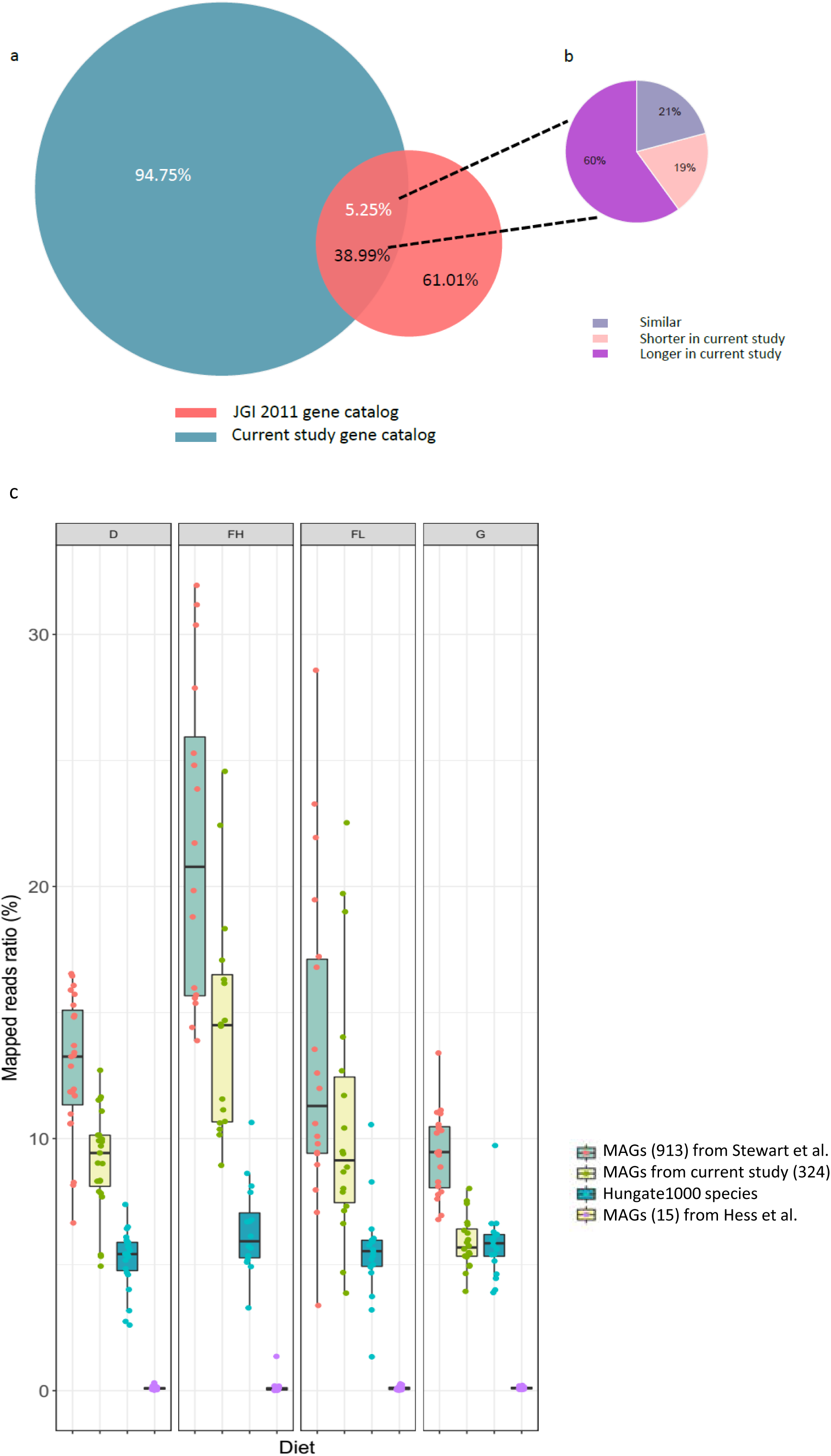
(a) Identity of current study genes compared to Hess et al. [14]. (b) Differences in gene length between studies. Green, the length of genes is longer in the current study; Blue, the length of genes is similar; Red, the length of genes is shorter in the current study. (c) Percentage of total reads in diet groups that mapped to MAGs. Mapping ratios of 77 samples to 913 genomes were calculated based on [6]. Mapping ratios of 77 samples to Hungate1000 isolates were calculated based on [5].

In order to compare our data with available genomes, MAGs were constructed based on abundance profiles and co-abundance clustering methods. We identified 324 MAGs with an average size of 1.8 Mbp (minimum threshold of 1 Mbp; see Methods for more information on these MAGs). More than half (173) were medium-quality and 23 were high-quality drafts (>90% completion, <5% contamination) [16]. Except one MAG annotated to the Archaea domain (Euryarchaeota), all were annotated to the Bacteria domain. For the bacterial MAGs, 39% could be annotated to the order level but only 2.5% (8 MAGs) to the genus level; all belonging to *Prevotella*. These rumen MAGs were compared to the Hungate1000 genomes [5] and to the 913 MAGs reported from Scottish cattle [6] Ninety-six (30%) out of 324 MAGs were similar to Scottish cattle MAGs but only 23 (7%) were similar to genomes from the Hungate1000 project, highlighting the novelty represented by draft genomes and the large diversity not yet covered in culture collections (Supplementary Table 3). In addition, we compared the proportion of mapping ratios of genomes and MAGs to an external dataset obtained from total rumen content samples of 77 cattle from two different genetic stocks that were fed diets characteristics of beef and milk production systems. Beef cattle, represented by the Charolais breed were fed fattening diets high (n=16) or low (n=18) in starch and lipids; whereas Holstein dairy cows were fed a corn silage and concentrate diet (n=23) or grazed a natural prairie (n=20) (Supplementary Table 4). The 324 MAGs were present in all four diet groups from our validation cohort; about 10% of reads from the 77 cattle dataset mapped to these MAGs. For genomes from the Hungate1000 project [5, 9], which are representative of the diversity of cultured rumen bacteria and archaea, the mapping rate was 5.4%, whereas the mapping rate for the largest MAGs collection of Stewart et al. [6] was around 13%. In contrast, only 0.1% of reads mapped to the 15 metagenomic species described by Hess et al. [14] (Figure 1c).

## Analyses

### Comparison of gastrointestinal microbiomes: bovine rumen versus Human, pig and mouse

Genes were taxonomically classified using CARMA3 [17] and compared to genes from the human, mouse and pig gut catalogs [11–13]. Up to 42.7% of rumen genes could be annotated to known phyla. This value is similar to pig gut (41.3%) but lower than the human (55.9%) and mouse gut metagenomes (59.6%) (Supplementary Figure 2 and Supplementary Table 5). Firmicutes and Bacteroidetes were predominant in all catalogs representing 84-94% of assigned genes and in accord with the expected gastrointestinal-associated microbial communities in mammals. For the rumen, however, the proportion of Firmicutes and that of Bacteroidetes was lower and higher, respectively, than for the other three catalogs. Other enriched phyla (>2%) in the rumen catalog were the Spirochaetes, Proteobacteria, Euryarchaeota, Actinobacteria and Fibrobacteres that, with the exception of Proteobacteria and Actinobacteria in Human, were more abundant in the rumen than in the other catalogs (Supplementary Figure 3). At the genus level, only 8.7% of rumen genes could be annotated; a value similar to that of the other two animal catalogs but lower than that of human (16.8%), reflecting a more extensive characterization of human-associated microbes. However, the top 10 enriched genera in the rumen showed distinct abundance patterns compared with the same genera in other catalogs (Supplementary Figure 4 and Supplementary Table 5 & 6). These differences in symbiotic microbial genera likely reflect dissimilarities in dietary lifestyles, anatomical localization of the gut fermentation compartment and are indicative of predominant functions, i.e. methane production and plant fiber degradation for ruminants.

*Prevotella* was the most abundant rumen genus with 39% of genus-annotated genes assigned. Other abundant genera were *Treponema*, *Butirivibrio*, *Methanobrevibacter* and *Ruminococcus* that were absent or at lower proportions in other catalogs, particularly in the human catalog.

### Carbohydrate active enzymes in the bovine rumen metagenome

The efficient deconstruction of structural plant polysaccharides by symbiotic gastrointestinal microbes is what sets ruminants apart from other livestock species. We have therefore analyzed carbohydrate active enzymes (CAZymes) in the rumen ecosystem to obtain insights into this important function for nutrition and health of cattle.

Glycoside hydrolases (GHs) and polysaccharide lyases (PLs) are the most relevant classes of CAZymes as they orchestrate the breakdown of plant material and of diverse polysaccharides which are encountered in the rumen ecosystem, i.e. host, fungal, and bacterial glycans. GHs and PLs are classified into sequence-based families (145 GH and 26 PL familes; [18] that display a pronounced specificity for a glycan category, thereby offering a functional readout of the degradative power of an ecosystem. The rumen catalog reported here encodes 545,334 CAZymes of which ∼290,000 have degradative activity that are affiliated to 114 distinct GHs families (97.4%) and 18 PLs families (2.6%). These 545,334 CAZymes were compared to GenBank and to the most complete dataset of assembled genomes from rumen samples available [6] (Supplementary Figure 5). Stewart and co-workers [6] described 69,678 CAZymes with 65% to 72% identity to other datasets, and 91% of novel CAZymes (defined as having <95% identity to other datasets). Our catalog displays similar features with average 73% identity to Genbank and Stewart’s MAGs and 89% novel CAZymes (482,759 sequences with <95% identity). This expands the size of the CAZyme set of Stewart et al. by almost 8 times and represents the most extensive source of reference CAZyme sequences in the rumen niche so far. It is noted that 32,755 degradative CAZymes are present in the Hungate1000 reference genomes [5].

In the rumen catalog, the substrate specificity of the most abundant GH families reflects the prominent glycan sources of herbivores: starch (GH13, GH77 and GH97, by decreasing abundance), pectins and hemicelluloses (GH43, GH28, GH10, GH51, GH9 and GH78, by decreasing abundance). In contrast, only one of the 15 most abundant families, namely GH25 lysozymes, targets a non-plant substrate (peptidoglycan). Additionally, three of the five most abundant families (GH3, GH2 and GH5) represent enzymes active on a wide range of substrates, not necessarily from plant origin. Two of these families (GH2 and GH3) contain exo-glycosidases that act on the oligosaccharides produced by depolymerases, a broad function that may explain their abundance.

Dockerins domains (DOCs) are key building blocks of cellulosomes and amylosomes complexes [19, 20]. The DOC sequences are found in modular proteins and help the protein to which they are appended to bind cohesin domains (COHs) found as repeats in large proteins named scaffoldins. This system allows the spatial grouping of numerous binding and enzymatic modules into large assemblies for a synergistic action of their components in the immediate vicinity of the bacterial cell. In the rumen catalog, more than 12,000 dockerin modules were identified. Intriguingly, some proteins harbored many dockerin modules, up to 13 modules in a single sequence, without any other recognizable functional module. The function of such polydockerin proteins is unknown, and polydockerin proteins were not observed in reconstructed MAG (max. of two DOCs in a protein). In the literature, dockerin modules initially detected in cellulosomes, have been investigated in relation to their co-occurrence with CAZymes in these cellulosome complexes [21, 22]. Surprisingly, our analysis of the rumen catalog reveals that only ∼24% of the DOC-containing proteins carry a CAZyme domain. The remaining DOC-containing proteins were subjected to a Pfam domain annotation, which identified proteases (4%) and some lipases (<0.3%), while a third of DOC-containing proteins are attached to non-catalytic modules, likely involved in the binding of these non-carbohydrate substrates. More importantly, the last third did not have any match to any Pfam domain (Supplementary Figure 6).

The CAZyme profile in the rumen catalog was compared to the mouse, pig and human reference gut catalogs [11–13]. Despite important differences in the size of these catalogs, similar trends could be observed on, for example, the ratio of DOCs or GHs plus PLs over the catalog size, or the most abundant GH families (Supplementary Table 7). The number of distinct GHs/PLs is also very similar, and a detailed analysis highlighted 101 GHs families common to all four catalogs, while only five GH families were specific to a single catalog (Supplementary Figure 7). These specific families were closely related to the hosts’ diets. In accord with herbivory, 305 GH45 cellulase modules were found in the rumen catalog against none in the human and mouse catalogs, and only 12 for the pig. In contrast, we identified families GH70 and GH68, transglycosidases acting on sucrose, and GH47, processing N-glycan, that are absent in the rumen but present in other catalogs. For instance, 94, 24 and 6 GH70 modules were found in the human, pig and mouse catalogs, respectively, whereas the rumen had zero occurrence.

The specific adaptation of the rumen microbiota to herbivory was confirmed by comparing its GH+PL family counts against the human catalog after normalization (Figure 2). The most enriched GH families in the rumen are involved in the degradation of plant polysaccharides while the more depleted families of GHs are those degrading animal (host) glycans. These observations are not only in accord with the normal diet of cattle normal diet but they are also in agreement with the absence of a glycoprotein-rich mucus lining of the rumen as opposed to the lower gastrointestinal tract. Finally, we also observed that multiple DOC module duplications seem to be more frequent and intense in the rumen as up to 13 DOC repeats in a single protein were found for the rumen catalog, compared to only six in the human, four in the pig and two in the mouse catalogs.

**Figure 2.**
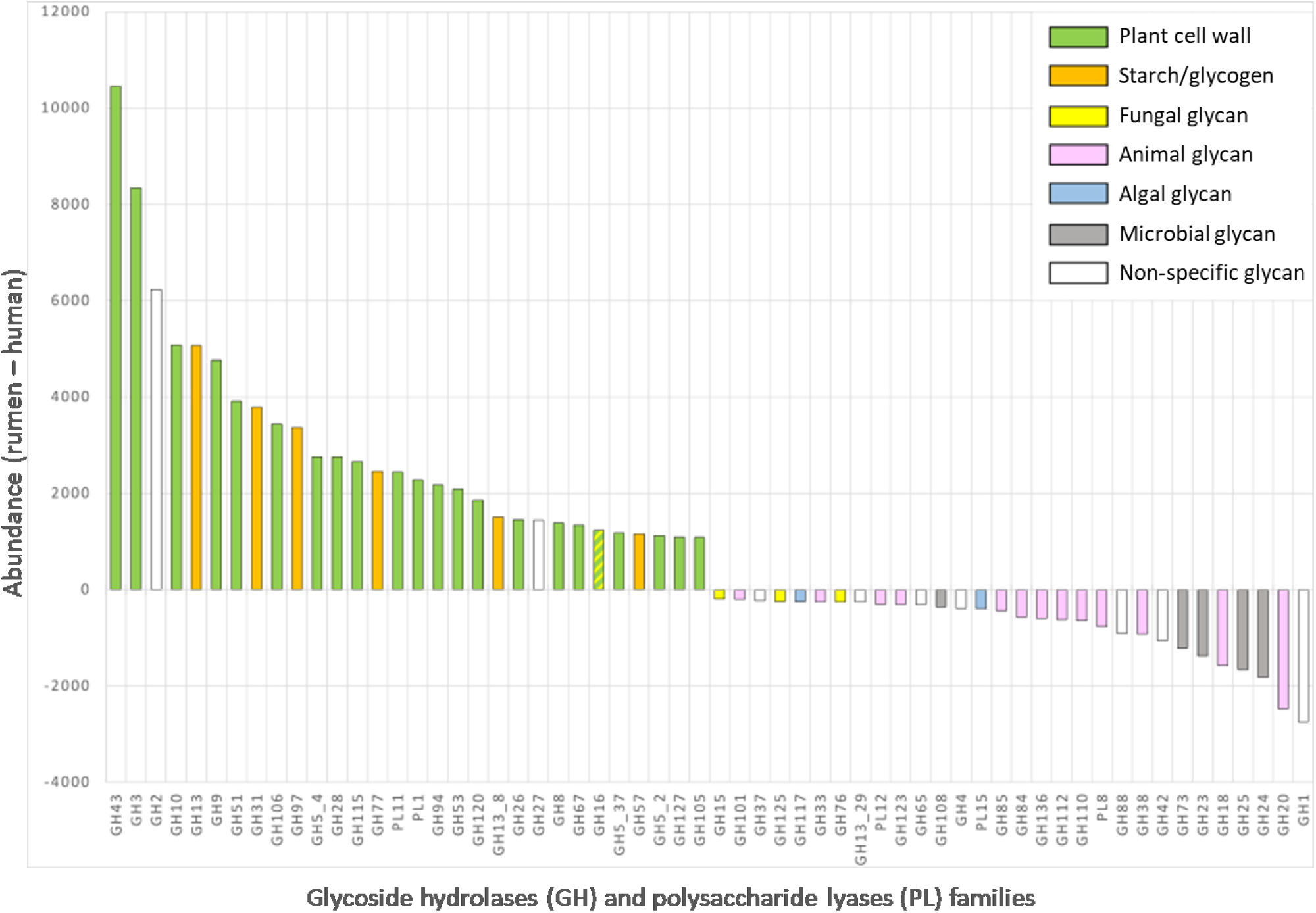
Enrichment or depletion of glycoside hydrolases and polysaccharide lyases in the bovine rumen as compared to human gut. Human counts were normalized to rumen catalog size before comparison.

CAZyme-encoding genes were also annotated in the 324 MAGs. Remarkably the most abundant families in the MAGs are for plant cell wall breakdown and correspond closely to the most abundant families in the non-redundant catalog. The CAZyme profiles of each generated MAG were thus determined and subjected to a hierarchical clustering analysis (Supplementary Figure 8) which that revealed that the MAGs roughly group together according to their predicted taxonomy, even despite large differences in repertoire size within each phylum. Hereafter, we analyzed in detail several strategies for carbohydrate foraging that have evolved in the different bacterial phyla. Among the predicted Firmicutes, MAGs encoding cellulosomes and amylosomes displayed a readily recognizable profile characterized by the presence of several DOC and COH domains along with several GHs families containing cellulases (GH5, GH44, GH48, and GH124) and amylases (GH13 with associated CBM26), respectively. We also identified Bacteroidetes MAGs that contained a few DOC domains but, interestingly, none of these MAGs contained a recognizable COH domain. The presence of dockerin domains not associated to cohesins in Bacteroidetes MAGs was recently reported in the moose rumen microbiome [7]. The role of the dockerins in Bacteroidetes is unclear but the conspicuous absence of cohesins suggests that they may not be needed for the assembly of a *bona fide* cellulosome or that the Bacteroidetes cohesins are so distantly related from their clostridial counterparts that they cannot be recognized.

Confirming previous reports in the literature [23], the largest CAZyme repertoires dedicated to plant degradation were found among the predicted Bacteroidetes members, which represent the majority of the 324 reconstructed genomes. In Bacteroidetes, CAZymes are often grouped in distinct Polysaccharide Utilization Loci (PULs) around *susC* and *susD* marker genes to build up specific depolymerization machineries capable of deconstructing in a synergistic manner even the most complex polysaccharides [24, 25]. In this context, it is interesting to note the clustering of families GH137 to GH143 recently shown to catalyze the breakdown of type II rhamnogalacturonan [24] in the CAZyme profile heatmap (Supplementary Figure 8). Inspection of the predicted PULs in the Bacteroidetes MAGs revealed the presence of degradation machineries dedicated to pectin (type II rhamnogalacturonan), starch, or barley β-glucan (Supplementary Figure 9).

Other MAGs with distinctive CAZymes were those assigned to Proteobacteria and Fibrobacteres that despite their small number (eight and six respectively) form tight groups. Predicted Proteobacteria were characterized by the presence of families GH84 and GH103 along with an important diversity of GH13 subfamilies. In contrast, the Fibrobacteres show the presence of several families known to degrade cellulose and β-glucans (*e.g.* GH5, GH45, and GH55). Focusing on CAZymes from *Fibrobacter* spp. present in the catalogue revealed an astonishingly strain-level diversity for this genus. We compared the CAZymes present in *Fibrobacter succinogenes* type species [26] against all Fibrobacter CAZymes in the catalog. There were 1262 hits with ≥ 90% identity to 135 of the 175 *Fibrobacter succinogenes* CAZymes, whereas only 19 of them had a 100% identity with the type strain. Up to 465 and 375 of these genes were differentially abundant in the Holstein and Charolais groups, respectively (Supplementary Table 8). Zooming in on a particularly important endoglucanase enzyme, GH45, reveals its presence in all 77 animals receiving different diets. Animals harbored between four to 13 GH45 variants and each gene was present in 25 to up 99% of all animals; however, the type strain, at 79%, was not the variant most commonly present.

### Common functions and influence of diet on the bovine rumen microbiome

To investigate how different feeds affected the rumen microbiota in beef and dairy cattle we examined samples from 77 cattle described above. By using this 77-sample dataset, differences in α-diversity were observed between diets at the gene level. Animals fed fresh grass had the highest α-diversity and richness compared to other diets containing conserved feeds. Particularly, animals on fattening diets had a lower α-diversity. In contrast, the fattening diet rich in starch and polyunsaturated fatty acids (PUFA) exhibited the highest β-diversity and/or had the highest disparity in interquartile range (box in the boxplot) for all indices (Figure 3). The rumen microbiome of animals fed this diet also exhibited the highest dispersion on ordination analyses at the gene level (Supplementary Figure 10). Such changes, akin to the described Anna Karenina principle [27] for microbiomes, probably reflected divergences in individual microbiomes (and hosts) responses to PUFAs and may underlie a stress response to the diet.

**Figure 3.**
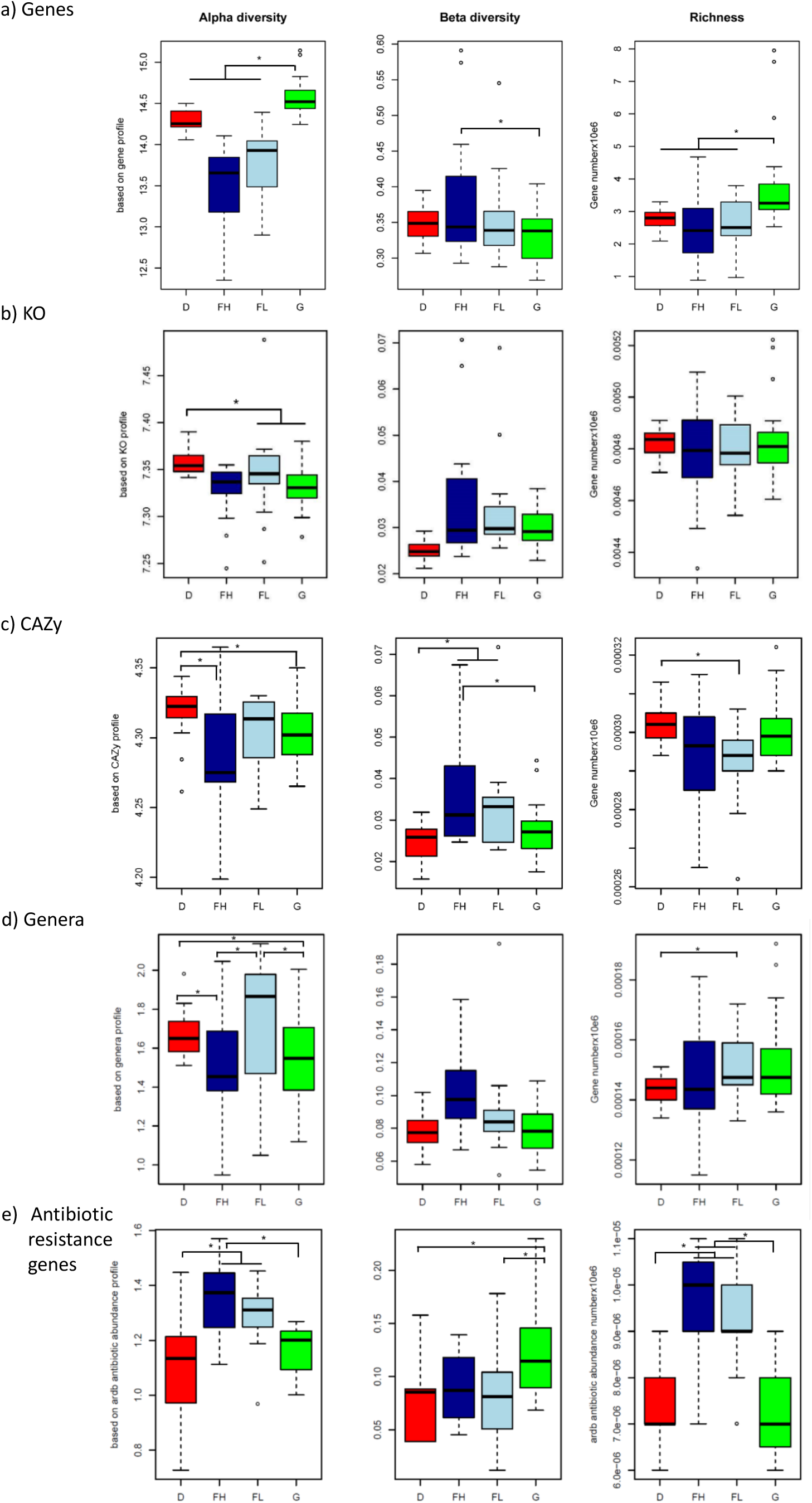
Effect of diet on diversity indexes of the bovine rumen microbiome. Comparison of Alpha diversity, Beta diversity and Richness at Gene (A), KO (B), CAZy (C), Genera (D) and antibiotic resistance gene (E) levels among cattle fed: dairy (red, n=23), fattening high-start (dark blue, n=16) fattening low-starch (light blue, n=18) and grazing (green, n=20) diets. * indicates P<0.05.

Genes were annotated to known functions (KEGG and CAZy) and taxonomical information was derived. For functions, there were 43.3% of the genes that could be classified into KEGG orthology and 2.1% assigned to feed carbohydrate degradation. A total of 5,893 unique KEGG orthologs (KOs) and 45,683 unique CAZy enzymes and binding modules were identified. Comparing the annotated genes for KEGG and CAZy functions showed a large overlap among groups with 91% and 94% of shared genes, respectively (Supplementary Figure 11). Contrasting with results on overall gene abundance, the highest α-diversity was observed for the corn silage diet group (Figure 3). To assess the functions encoded by the minimal rumen metagenome, we identified genes and KOs that were shared by all individuals in the group of 77 cattle. We found common sets of non-redundant genes, functions, genera and MAGs that were shared by all 77 rumen samples (Figure 4 and Supplementary Figure 12). The core gene set shared by all animals represented only <0.1% (6051 to 12075 genes depending on the calculation method–see Methods) of the nearly 14 M non-redundant genes in the catalog, whereas about 63% of the KO functions (∼3700) were shared indicating the high redundancy of genes for similar functions. Compared to all annotated KO, this minimal KO set was significantly enriched in pathways related to metabolism (amino acids, carbohydrate, nucleotides and metabolism of cofactors and vitamins), cellular processes (motility), and genetic information processing (translation) (Supplementary Figure 12b). Concerning the diversity of genera found in the different groups, there was also a relatively large overlap. Out of 242 genera identified by the taxonomic analysis described above, 182 (75%) were present in all four groups but only 67 (27%) were shared by all animals (Figure 4 and Supplementary Figure 11). This overlap was maximal for MAGs identified in this study, which were present in virtually all individuals (Figure 4). The presence of common functions may explain the plasticity of the microbiota and adaptability of ruminants to digest various types of feeds even after sudden dietary changes. To get a better understanding of the functional changes induced by diet in these microbial communities, we analyzed the abundance of genes in the 77-sample dataset for functions, genera and MAGs. To avoid possible confounding effect of breed and sex, the differential abundance analysis was performed within each breed. For Holstein, greater changes in the relative abundance of genes were observed; ∼43% difference in KEGG and CAZy functions (Supplementary Tables 9 and 10, and Supplementary Figure 13 a & b). For CAZy, 146 catabolic families exhibited indeed differences in abundance between the corn silage and grazing groups (Supplementary Figures 13a and 14, Supplementary Table 10). Most of the differences related to functions were due to increases in the relative abundance of genes in cows fed the corn silage diet rather than the presence of different genes.

**Figure 4.**
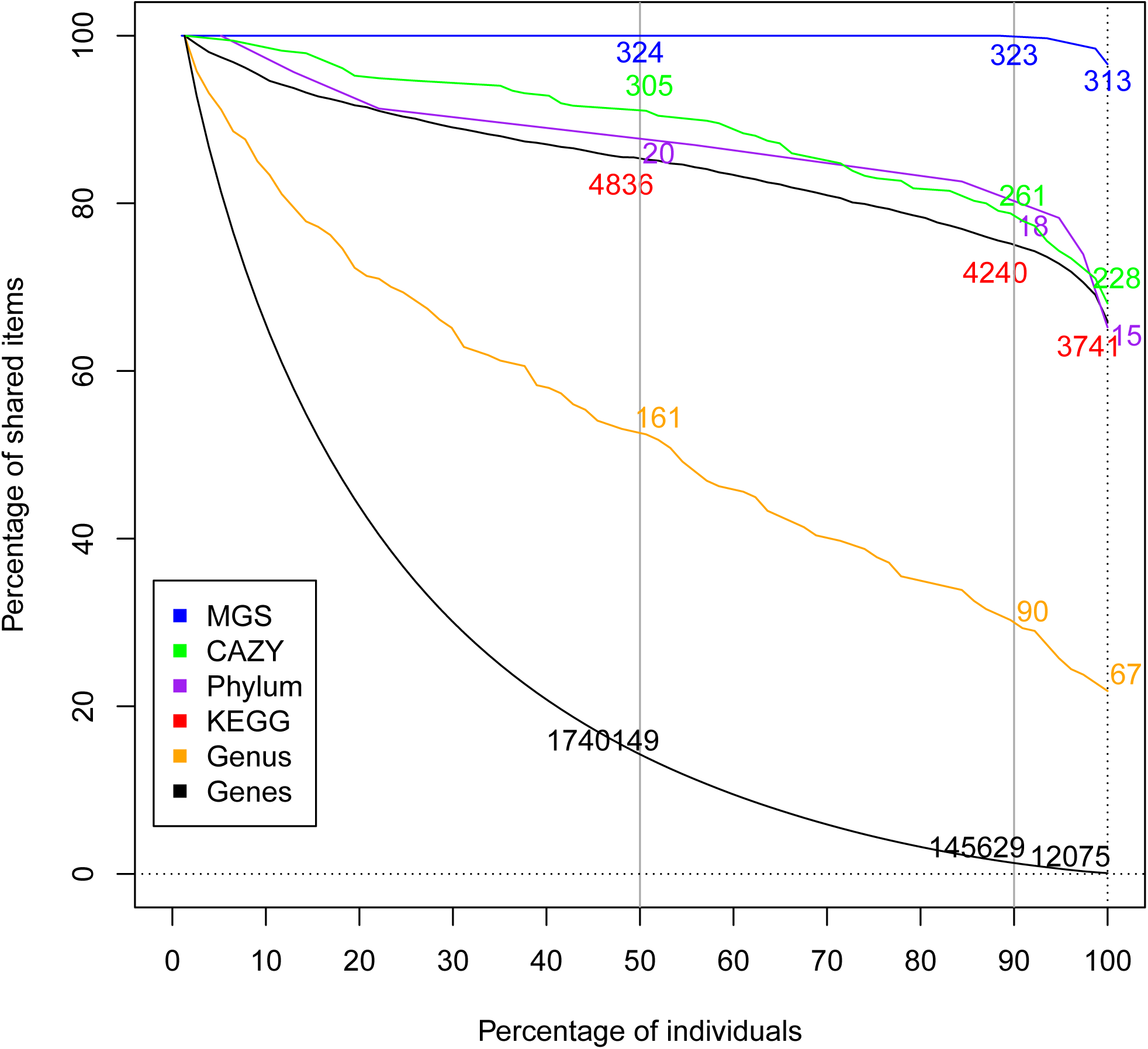
Size of the shared microbiome features among cattle (n= 77) fed four different diets for the number of genes (black), genera (orange), phyla (purple), MAGs (blue), KEGG pathways (red), and CAZy (green). The percentages of shared items and animals are represented on the y and x axes, respectively. The absolute numbers for each item are indicated at the intercept between the percentages of items and animals at the thresholds of 50, 90 and 100%. Only ∼1% of genes were shared by 90% of the cattle, whereas close to 80% of KO and CAZy functions were shared by 90% of the cattle, suggesting gene redundancy for similar functions. To note the presence of most MAGs assembled in this work in 90% of the cattle.

Notwithstanding, the greatest contrast was observed for families targeting fructans and sucrose that were more abundant in the grazing group. Particularly for family GH32 (P = 7.6E-12) whose higher abundance could be related to the high contents of sucrose and fructans in grasses [28, 29] included in the grazing diet. The other CAZy families differing in abundance were all more abundant in the corn silage-diet group. Interestingly, these results highlight the ability of ruminal bacteria to be equally capable of using glycans from plants as well as from microbial origin such as bacterial peptidoglycans, bacterial exopolysaccharides and fungal cell walls. Corn silage, the main constituent of the diet, is a fermented feed with an abundant epiphytic microbiota composed of exopolysaccharide-producing lactic acid bacteria, fungi and yeasts [30, 31]. Accordingly, the CAZome of the corn silage-diet group was oriented towards degradation of starch, a nutrient abundant in corn silage and practically absent in the grazing diet. Forty-two CAZy families targeting plant cell wall polysaccharides were also overabundant in the corn silage-diet group. This could reflect the diversity of fiber structures that ruminal bacteria have to face when cows are fed with such a diversified diet in terms of plant fractions and botanical origins (whole corn plant and soybean meal in the corn silage diet against a natural prairie, composed predominantly of grasses in the grazing diet). Finally, the overabundance of CAZy families targeting animal glycans in the silage-fed cohort was striking since no glycoprotein-rich mucus is secreted in the bovine rumen as opposed to the lower gastrointestinal tract [32]. It is possible that this difference reflects that CAZy families targeting animal glycans harbor numerous enzymes that are not fully characterized and may be able to act on plant or even fungal glycans, which contain a panel of osidic constituents that are very similar to that of animal glycans. Enzyme promiscuity may indeed confer metabolic flexibility and an ecological advantage to certain microbes in the gut ecosystem.

For genera and MAGs, up to 44% (106 genera) and 58% (188 MAGs) of the total detected were differently abundant in the microbial communities of the two cows’ groups (Supplementary Table 11 and Supplementary Figure 13 c & d). *Fibrobacter* and *Ruminoccocus* were more abundant in the corn silage diet group whereas *Prevotella*, *Butyrivibrio* and *Methanobrevibacter* were more abundant in the grazing group.

For Charolais on fattening diets differing in starch content, less than 5% differences were observed in the abundance of genes for functions or genera. Only eight CAZy families exhibited differences in abundance between the two Charolais groups, the differences in abundances being less significant than for the Holstein groups (p = 0.004) (Supplementary Figures 13a and 14, Supplementary Table 10). The absence of marked variations in the abundance of glycoside-degrading enzymes between the two fattening diets reflects indeed their similar composition. The differences in starch content were not great enough to drastically impact the carbohydrate harvesting functions of the ruminal microbiota, at least at the gene level. Similarly, smaller differences in the abundance of genera and MAGs were detected between these two diets (Supplementary Table 11, Supplementary Figure 13 c & d).

Metadata collected on the Holstein and Charolais animals were analyzed using a vector fitting method on the top of the bidimensional NMDS ordination (Supplementary Table 12). Diet had a significant effect on metagenome gene distribution, particularly in the Holstein group (r^2^= 0.68, P = 0.0001), but also variables such as live weight, feed intake, and rumen volatile fatty acids were significant. Protozoal numbers were also a significant variable explaining the distribution of genes in the metagenome of animals, underpinning their importance as key members of the rumen ecosystem and modulators of the prokaryotic community.

### Antibiotic resistance genes

The spread of antibiotic-resistance pathogens in the environment is a great concern in public health. Livestock species are a known reservoir of antibiotic resistance genes (ARG) [33]. Information from ruminants is predominantly from the fecal microbiome [34], and although the importance of the rumen microbiome has also been highlighted [35, 36], data on the rumen resistome is still fragmented. As an example of the useful information than can be retrieved from a gene catalog, we evaluated the presence of ARG in the rumen microbiome as previously reported [12]. Forty-two ARGs encoding resistance to 27 antibiotics were detected in the catalog. The most abundant resistances were to tetracycline and bacitracin with Charolais animals harboring globally a higher proportion of these genes (Supplementary Figure 15), probably reflecting the effect of diet [35]. It is noted that antibiotics as growth promoters were never used on these animals. In both the bovine rumen and the porcine gut [12], the most abundant ARGs confer resistance to tetracycline and bacitracin. The diversity of ARG is low compared to pig feces where resistance to up to 52 antibiotics was reported, even in farms with no use of growth promoting antibiotics [12]. Similar to this study, tetracycline resistance was reported as highly abundant in the rumen, otherwise prevalence of resistance to other antibiotics varies between studies [36, 37]. Although the methodologies used to detect ARGs could play a role in these differences [35–37], it is probable that variation in the rumen resistome may differ between countries and regions as it can reflect decades of exposure since antibiotics started to be used in farms.

## Discussion

Ruminants are extraordinary bioreactors, engineered by nature to use recalcitrant plant biomass—a renewable resource— as feedstock for growth and for production of useful products. This ability is a microbial attribute that was important in domestication and that today has a renewed interest due to human population increases, resource scarcity, and climate change issues. The reference gene catalog from the rumen microbiota reported here is a useful resource for future metagenomics studies to decipher the functions and interactions of this complex ecosystem with feeds and the host animal. Comparison with human, mouse and pig gut catalogs shows the distinct character and potential of the rumen ecosystem. As opposed to the microbiome of single-stomached animals including humans, the rumen microbiome harbors a plethora of genes coding for glycoside hydrolases (CAZymes) that degrade structural polysaccharides. Information on these enzymes that deconstruct biomass plant material and are essential for transforming recalcitrant feeds into meat and milk is also useful for the design of improved processes for the biofuel industry [38, 39].

The type of diet modulated as expected the abundance of genes and the metagenome profile of individuals. However, more than 90% of genes coding for functions (KO and CAZy) were shared among animals receiving different diets. This large functional diversity might be the key that allows ruminants to feed on a variety of dietary sources and to adapt to seasonal or production-imposed dietary changes. The 13.8M genes catalog produced in this work, despite being significantly larger than gut bacterial catalogs from other species [11–13] does only partially cover the diversity present in the rumen microbiome indicating the higher complexity of this ecosystem. The catalog needs to be expanded with additional data, particularly the inclusion of ciliated protozoa and fungi to reflect the overall diversity. Nevertheless, this catalog and the 324 uncultured assembled genomes are an important instrument to characterize and understand the biological functions of the rumen microbiome. This information is essential to enhance the sustainability of ruminant production.

### Methods

This study was conducted using the animal facilities at the French National Institute for Agricultural Research (INRA) in Theix and Bourges, France. Procedures on animals used in this study complied with the guidelines for animal research of the French Ministry of Agriculture and all other applicable National and European guidelines and regulations.

#### Rumen Sampling

Total rumen content samples from 10 animals used for deep sequencing metagenome were taken at the experimental slaughterhouse of the INRA Centre Auvergne-Rhône-Alpes. Total rumen content samples from 77 animals were also collected. These 77 animals, from two different genetic stocks, were fed diets characteristics of beef and milk production systems. Beef cattle, represented by Charolais breed were fed fattening diets high (n=16) or low (n=18) in starch and lipids; whereas Holstein dairy cows were fed a corn silage and concentrate diet (n=23) or grazed a natural prairie (n=20) (Supplementary Table 13). Rumen samples from these animals were also collected at the experimental slaughterhouse except for the grazing group. Cows from this latter group were fitted with rumen cannula and samples were taken from live animals.

#### Sample handling and DNA extraction

The 10 rumen samples used for deep sequencing were depleted from eukaryotes using washing and centrifugation steps. Rumen contents were filtered through a 400 µm nylon monofilament mesh. The filtrate was centrifuged at 300 *g*, 5 min to decant protozoa and the supernatant (fraction A) was stored at 4 °C. Fifty grams from the filtered rumen content retentate were mixed with 100 ml of anaerobic phosphate saline buffer (PBS), mixed manually for 5 min by gentle inversion, centrifuged at 300 *g*, 5 min to decant protozoa and the supernatant, passed through a 100 µm filter (fraction B), was stored at 4 °C. The pellet was mixed with 75 ml anaerobic, ice-chilled 0.15% Tween 80 in PBS and incubated on ice for 2.5 h to detach microbes attached to feed particles. At the end of the incubation, contents were vortexed for 15 s and centrifuged at 500 *g*, 15 min. The supernatant (fraction C) was mixed with fraction B and 50 ml of fraction A and centrifuged at 20,000 *g*, 20 min, 4 °C. The supernatant was decanted and the microbial pellet was exposed to an osmotic shock to lyse any remaining eukaryote (mainly protozoal) cells followed by an endonuclease treatment. Briefly, the pellet was suspended in water (Millipore Waters Milli Q purification unit) and incubated for 1 h at room temperature followed by DNase treatment (Benzonase, Novagen) as described [40]. The suspension was filtered through a 10 µm monofilament textile, collected by centrifugation as before, suspended in an appropriate volume of PBS and stored at −80 °C until DNA extraction. DNA was extracted following the method described by Yu and Morrison [41]. Samples from 77 animals were extracted directly from whole rumen contents using the same extraction method.

#### DNA library construction and sequencing

Paired-end (PE) metagenomic libraries were constructed and sequenced following Illumina HiSeq2000’s instruction. Quality control and bovine DNA removal (by aligning reads to *Bos taurus* genome Btau_4.0 [42]) for each sample were independently processed using MOCAT pipeline as previously described [10]. On average, 111.3 Gb of high-quality reads were generated for each of the 10 deep sequencing samples and 3.43 Gb (median ∼2.5 Gb) for each of the 77 samples (Supplementary Table 4). The averaged proportion of high-quality reads among all raw reads from each sample was 92.29%.

#### Public data use

The four public rumen microbial datasets used in this study include: (i) a cow rumen microbiome sequenced at DOE’s Joint Genome Institute (JGI) in 2011 (JGI 2011), which consists of 268 Gb of metagenomics sequences, 2,547,270 predicted genes and 15 uncultured microbial genomes assembled from the cow rumen [14] (NCBI accession number SRA023560), (ii) 8 rumen metagenomics samples from beef steers [15] (European Bioinformatics Institute (EBI), PRJEB10338), (iii) 501 rumen microbial genomes from the Hungate1000 Project (Integrated Microbial Genomes (IMG), JGI Proposal Id: 612 / 300816) and (iv) 913 draft microbial genomes assembled from Scottish cows’ rumen (EBI, PRJEB21624).

Three public gut microbial gene datasets from human^1^ (GigaDB, doi:<u>10.5524/100064</u>), mouse [13] (GigaDB, doi:10.5524/100114) and pig [12] (EBI, PRJEB11755) were also collected.

#### Construction of the rumen microbial gene catalog

High quality reads from 10 deep sequenced samples were processed in MOCAT toolkit [10] including de novo individual assembly (SOAPdenovo v1.06 [43], -K 47). The assembled contigs with length equal to or greater than 100 bp were used for gene prediction (MetaGeneMark [44], –M 100 –A) and redundant genes were removed (CD-HIT [45], ≥ 95% identity and ≥ 90% overlap), resulting in a non-redundant rumen microbial gene catalog containing 13,825,880 genes (Supplementary Table 1).

#### Evaluation of current rumen microbial gene catalog

To assess the representative of our rumen gene catalog, we used the largest rumen gene catalog published to date by Hess, et al.[14]. First, the genes with gaps were filtered as follows: genes were broken when meet ‘*N*’ base, a subset for each interrupted gene was obtained, retaining only the longest sub-gene as representative of the original gene. A total of M genes without gaps were obtained, termed ‘JGI-2011-gene-catalog’ and used for following analysis (Supplementary Table 2).

Further, 13.83 M genes from current study and 2.46 M genes from JGI were pooled together to identify shared genes using CD-HIT [45]. The comparison of gene length between the two catalogs was conducted as previously described [11].

#### Gene catalog annotation

Taxonomic assignments of genes from rumen, mouse, pig and human guts were performed using CARMA3[17] on the basis of BLASTP [46] (V2.2.24) against the NCBI-NR database (v20130906 for rumen, mouse, pig guts; v20160219 for human gut) (Supplementary Table 5). Microbiotas from these four species were compared at different taxonomic levels. Functional annotation based on Kyoto Encyclopedia of Genes and Genomes (KEGG) database was performed using an in-house pipeline [11]. Annotation of the carbohydrate-active enzymes (CAZymes) of each catalog was performed by comparing the predicted protein sequences to those in the CAZy database and to Hidden Markov models (HMMs) built from each CAZy family [47], following a procedure previously described for other metagenomics analyses [7]. In order to allow a direct comparison of the results, annotation of antibiotic resistance genes (ARGs) was done as previously reported in the pig metagenome catalogue [12] by using the ARDB database [48].

#### Construction relative abundance profiles of genes, KOs, ARG and CAZY enzymes

The gene profiles of 77 rumen samples were generated by aligning high-quality clean reads to the current 13.83M gene catalogue (SOAP2, ≥95% identity) [49]. Gene relative abundance was estimated as described previously [50]. The relative abundance of each KEGG orthologous group (KO), ARGs and CAZy enzyme was calculated from the abundance of its genes [11].

#### Characterization of total and minimal metagenome

We computed the total and shared number of genes, KO and CAZy functions present in random combinations of *n* individuals (with n=2 to 77, 100 replicates per bin) [50]. Furthermore, we used a permutation test to identify the second-level KEGG functions that were significantly enriched or depleted in the minimal KO set compared with the total KO set. We first calculated the contribution of second-level functions using the following formula:

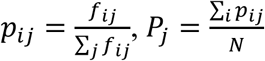

Where *f*_*ij*_ is the number of second level function *j* from the KO *i*; 𝑝_*ij*_ is the relative contribution of second level function *j* in the KO *i*; *N* is the number of KO in the KO set; 𝑃_*j*_ is the relative contribution of function *j* in the KO set.

Randomly sampling 999 times in all annotated KO set, simulated the distribution of each function. Calculating the position of this function contribution ratio of minimal KO set under the distribution of all annotated KO set. A *p* value of less than 0.01 was regarded as significant (Supplementary Figure 12b).

#### Construction of metagenomic species (MAG) and taxonomic assignment

To recover the draft bacterial and archaeal genomes from the 10 deep sequenced samples, we developed an in-house pipeline that comprises three steps as indicated in Supplementary Figure 16 and described below.

##### 1. Construction of Scaftig-Linkage Groups (SLGs)

We generated a scaftig abundance profile by aligning high-quality clean reads from 77 rumen samples to assembled scaftigs from samples [49]. Scaftig relative abundance was determined using the same method applied for gene abundance [49]. The highly co-abundance correlated scaftigs from each deep sequencing sample were binned into scaftig-linkage groups (SLGs) using the previously described pipeline [49] with modified parameters as follows, an edge was assigned between two scaftigs sharing Pearson correlation coefficient > 0.7 and the minimum edge density between a join was set as 0.99. A total of 745 preliminary SLGs with length > 1Mbp were generated for further analysis.

##### 2. Filtering of Preliminary SLGs based on GC content and assembly outputs

For all preliminary SLGs, we then examined their specificity by plotting the GC content versus reads aligned depth of each scaftig. In this step, 520 SLGs containing sole GC cluster were treated as ‘qualified’ and retained for the step 3. For the remaining 225 SLGs, 184 presented a scattered GC distribution and were discarded whereas the 41 SLGs containing two or more GC clusters were further processed. First, those SLGs with scaftig N50 <2000bp were considered as too fragmented and discarded. Then, multiple GC clusters in remaining SLGs were separated by DBSCAN[51] (Eps<=0.10, MinPts>=49). After splitting and filtering, we retained 55 ‘qualified’ SLGs that had a coverage depth greater than 20×.

##### 3. Reconstruction of metagenomic species

In order to improve the completeness and remove the redundancy of multiple metagenome assemblies from 10 deep sequencing samples, we performed hierarchical clustering for these 575 qualified SLGs based on their scaftigs nucleotide identity calculated by MUMi [52]. The MUMi distance between two SLGs (a and b) was defined as:

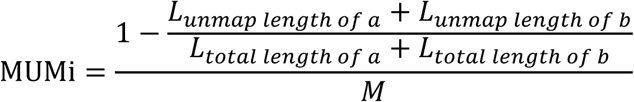

Where *M* = 2 × min(*L*_*total length of a*_, *L*_*total length of b*_)/*L*_(*total length of a*_ + *L*_*total length of b*_), *L*_*total length of a*_ is the length of SLG a, and *L*_(*unmap length of a*_ is the length of unmapped sequence compared with SLG b. The threshold for generating a species level metagenomic species (MAG) at 0.54 for MUMi as previously suggested [53]. Two hundred and eighteen qualified SLGs could not be clustered with other SLGs and were defined as singleton-MAGs. The remaining 357 qualified SLGs were clustered into 105 candidate-MAGs. We performed overlap-based assembling on the scaftigs for each of these 105 candidate-MAGs respectively, using Phrap with default parameters. To get reliable contiguous sequences for each candidate-MAG, the overlaps between two scaftigs less than 500bp were considered as unreliable and re-broken.

The 105 reconstructed candidate-MAGs were further examined using GC patterns using the same method mentioned in step 2 above. Eighty out of the 105 candidate-MAGs containing sole GC cluster were retained as combined-MAGs. The remaining 25 candidate-MAGs containing two or more clusters were split into sub-MAGs using the same method mentioned in step 2 above. In order to preserve the most comprehensive genomic information for these sub-MAGs, sequences from each sub-MAG was aligned back to its original SLGs. If the sub-MAG covered 90% or more sequences of its original SLG, it would be retained as a revised-MAG. Otherwise, its original SLG will replace the corresponding sub-MCs and be considered as a revised-MAG. This splitting step finally obtained 31 revised-MAGs.

After filtering the total sequence size of 218 singleton-MAGs, 80 combined-MAGs and 31 revised-MAGs with the criterion of > 1Mbp, we finally obtained 324 MAGs for rumen microbiota including 224 singleton-MAGs and 100 combined-MAGs (Supplementary Table 15). We used the same pipeline described above for the gene catalog for the ORF prediction and taxonomic annotation of MAG genes. We used CheckM [54] to estimate of the completeness, contamination and heterogeneity of metagenomic species (Supplementary Table 15). MAGs were assigned a taxonomic level annotation if more than 50% of its genes were assigned at a given taxonomic level (including genes with no match) (Supplementary Table 16). The MAG relative abundance of 77 rumen samples was calculated from the relative abundance of its aligned genes.

#### Quality assessment and taxonomic annotation of MAGs

CheckM software [54] was used to calculate the completeness and contamination of these MAGs. The median percentage of completeness was high at 62.5% with a low, 2.6% contamination. The combined-MAGs showed higher completeness but also slightly higher levels of contamination and strain heterogeneity than singleton-MAGs (Supplementary Figures 17 and 18). Taxonomic annotation for rumen MAGs was performed using CARMA3 on the basis of BLASTP against the NCBI-NR database (v20130906) and compared with MAGs from pig and mice (Supplementary Table 15, Supplementary Figure 19).

#### Cluster distribution by diet at species level

The relative MAG abundance profile (matrix of 324 × 77) obtained above was analyzed to highlight differences induced by diet. As we found when coverage of a MAG is less than 0.1 the depth of this MAG is close to 0 (Supplementary Figure 20). This result is caused by the noise and is non-conducive to the MAG clustering. Therefore, when the coverage value was less than this threshold value, then we set the value of depth equal to 0.

#### Ordination and differentially abundance analyses

Breed and diet distribution were visualized in ordination analyses based on two-dimensional non-metric multidimensional scaling [55]. Dissimilarity between pairs of samples was calculated using Bray–Curtis dissimilarity index [56]. Vegan R package [57] was also employed to estimate the diversity indexes corresponding to richness, alpha (Shannon index) and beta diversity (Whittaker). The ‘*envfit*’ function of Vegan was used to determine whether phenotype information corresponding to the 77 samples contribute to the overall pattern of the rumen microbiome structure. The significance of the environmental factors was assessed after 9999 permutations.

The relative abundance of the 13,825,880 non-redundant genes was collapsed into taxonomic (Phylum and Genus) and functional levels (KEEG and CAZy). Procrustes rotation analysis was performed to compare the ordinations obtained at different levels. Identified KOs were mapped to KEGG and visualized using the Interactive Pathway Explorer (iPath2.0) web-based tool [58]. To estimate a core, the overlapping number of Genus, CAZy and KOs between Holstein and Charolais breeds was compared.

To avoid confounding factors such as: sex, breed and age, the differentially abundance analysis was performed within breeds. Therefore, for each breed, diet comparison was done based on a Zero-Inflated Gaussian mixture model as implemented in the fitZig function of the metagenomeSeq R package [59]. Correction for multiple testing was done, and the cut-off of the differential abundance was set at FDR ≤ 0.05.

## Availability of supporting data and materials

Metagenomic sequencing data generated in this study have been deposited in EBI database under the accession code PRJEB23561. The data of assembled scaftigs, the rumen gene catalog, the rumen MAG catalog, and the abundance profile tables generated in this study have been deposited in *GigaScience Database* (DOI: 10.5524/100391).

## List of abbreviations

ARG: antibiotic resistance genes
bp: base pairs
CAZymes: carbohydrate active enzymes
COHs: cohesin domains
DOCs: dockerins domains GHs glycoside hydrolases
KEGG: Kyoto Encyclopedia on Genes and Genomes
KOs: KEGG orthologs
MAGs: metagenome-assembled genomes
ORF: open reading frames
PLs: polysaccharide lyases
PBS: phosphate saline buffer
PUFA: polyunsaturated fatty acids
PULs: polysaccharide utilization loci
SLGs: scaftig-linkage groups

## Declarations

### Ethics approval

Procedures with cattle were conducted in accordance with the guidelines for animal research of the French Ministry of Agriculture and applicable European guidelines and regulations for experimentation with animals (Certificate of Authorization to Experiment on Living Animals No. 004495 and ethics committee notification 10726-2016062616304407 V4)

### Consent for publication

Not applicable

### Authors’ contributions

JL, HZ, SDE, and DPM designed the work and managed the project. JL and HZ designed the analyses, and analyzed and interpreted the sequencing data. JL, HZ, SDE, GPV, BH, NT, VL, YRC, JE, and DPM wrote the manuscript. GPV, BH, NT, and VL conducted data analysis on CAZy and were involved in the interpretation of data. YRC and JE conducted integrative data analysis, implemented ARG analysis and were involved in the interpretation of data. MP and CM were involved in the implementation of animal studies, samples and metadata collection. ZY, HZ, ST, and FL performed data analyses, constructed and annotated the MAG catalog. WC, BC, and JLI performed data analyses, constructed and annotated the gene catalog. JG contributed to the experimental development and discussion for filtering rumen samples. MP, EM, XX, HY, LM, and JW contributed to text revision and discussion. KK interpreted the data, revised the paper. DPM coordinated the project. All authors approved the submitted versions and agree to be accountable for all aspects of the work.

## Acknowledgements

The authors acknowledge technical support for animal care, sampling and analytical measures on live animals provided by the personnel at INRA’s experimental units of Herbipôle (P. Faure and D. Roux) and Bourges, and the Herbivores research unit (Y. Rochette, F. Anglard, and B. Sepchat). Particular thanks to D. Graviou for DNA extraction and biochemical analysis.

We thank the direction of INRA divisions Microbiology and Food Chain, Animal Genetics, and Science for Food and Bioproduct Engineering for their support. We also thank E. Forano, and M. Naves (INRA) for helpful discussions during the setup of the project.

