## Supplementary material for "A catalog of microbial genes from the bovine rumen unveils a specialized and diverse biomass-degrading environment": Suppl. Fig & Tables

### FIGURES and TABLES

Supplementary Table 15. Excel file “Supplementary Table 14_Assessment of 317_MAG” 34
Supplementary Table 16. Excel file “Supplementary Table 16_rumen-pig-mice_MAG_annotation” ........ 34
Supplementary Table. Excel file “Supplementary Table 16_DA_KO” ........ 34

### Results

#### Constitution of a bovine rumen prokaryotic gene catalog

##### Supplementary Figure 1

a)

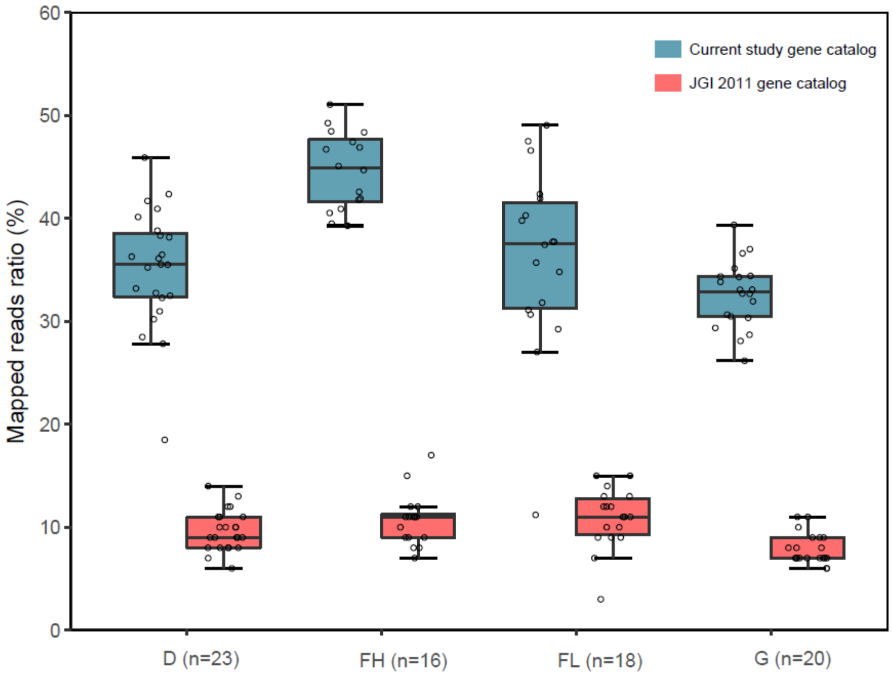

b)

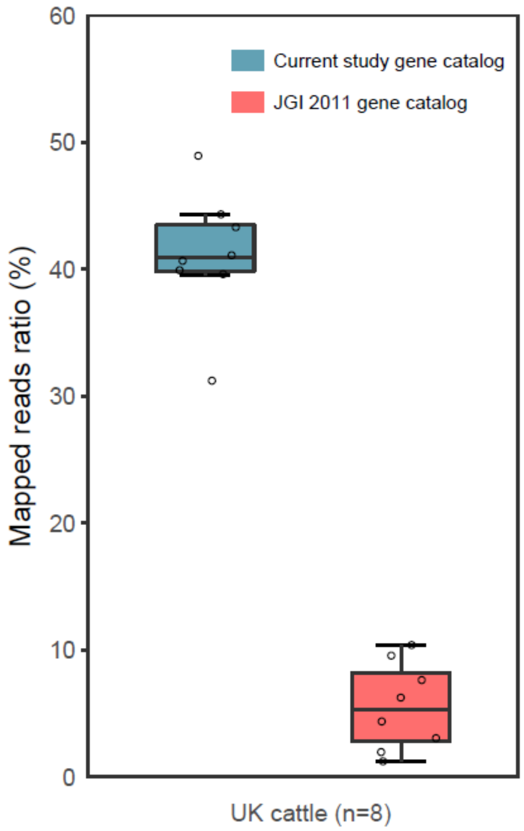

Supplementary Figure 1. (a) Percentage of total reads in the current study (n=77 samples) in four different diet groups that could be mapped to current study gene catalog and JGI 2011 gene catalog. (b) Percentage of total reads in unrelated studies of 8 UK cattle samples that could be mapped to the current study gene catalog and JGI 2011 gene catalog.

**Comparison of gastrointestinal microbiomes: bovine rumen versus Human, pig and mouse**

**Supplementary Figure 2**

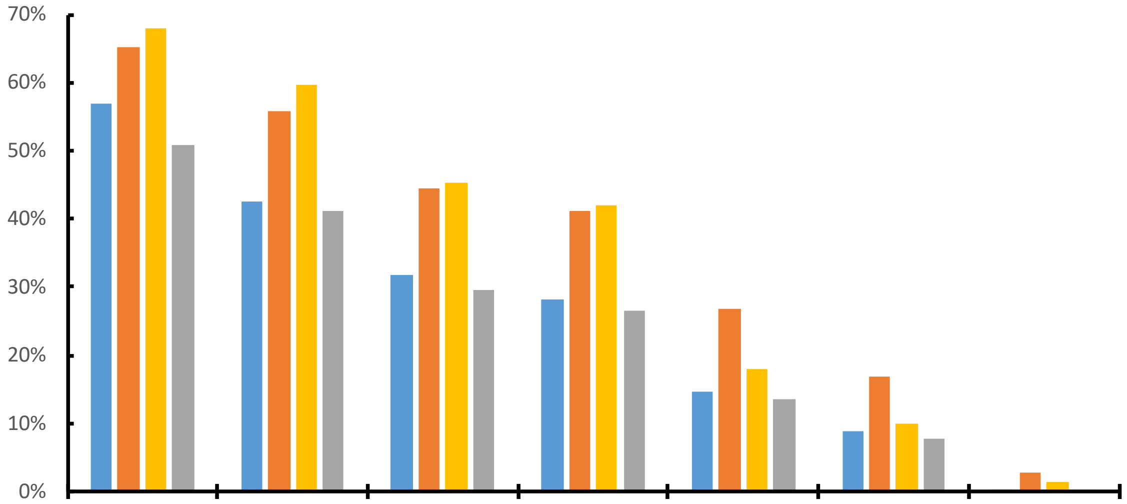

| | Superkingdom | Phylum | Class | Order | Family | Genus | Species |
|---|---|---|---|---|---|---|---|
| Rumen | 57.03% | 42.67% | 31.73% | 28.15% | 14.62% | 8.72% | 0.09% |
| Human | 65.27% | 55.94% | 44.44% | 41.10% | 26.69% | 16.79% | 2.79% |
| Mouse | 67.85% | 59.60% | 45.36% | 41.98% | 18.07% | 9.88% | 1.40% |
| Pig | 50.79% | 41.27% | 29.54% | 26.50% | 13.45% | 7.65% | 0.33% |

Supplementary Figure 2. The composition of four gene catalogs at different taxonomic annotation levels

**Supplementary Figure 3**

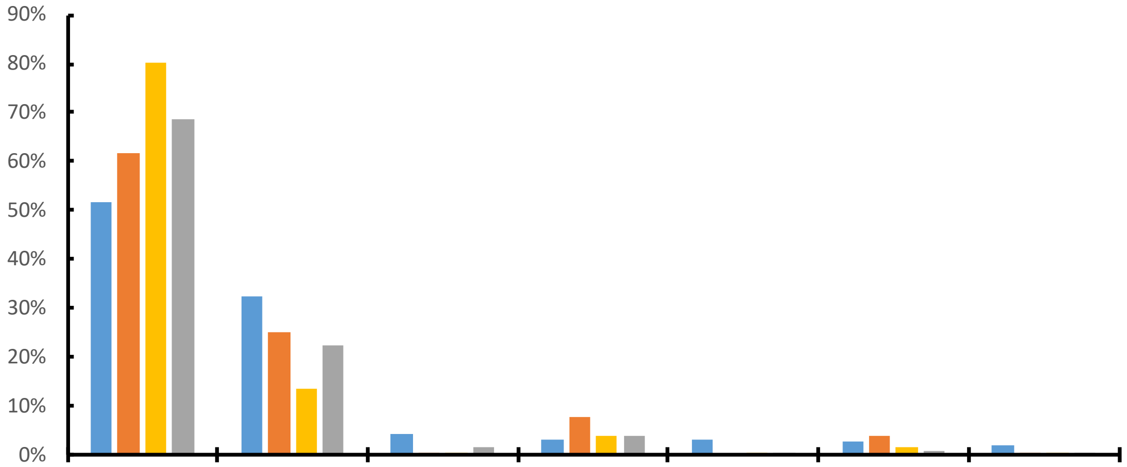

| | | Firmicutes | Bacteroidetes | Spirochaetes | Proteobacteria | Euryarchaeota | Actinobacteria | Fibrobacteres |
|---|---|---|---|---|---|---|---|---|
| Rumen | ■ | 51.67% | 32.31% | 4.19% | 3.06% | 2.90% | 2.59% | 2.01% |
| Human | ■ | 61.63% | 24.86% | 0.47% | 7.51% | 0.00% | 3.86% | 0.02% |
| Mouse | ■ | 80.18% | 13.56% | 0.10% | 3.99% | 0.03% | 1.36% | 0.02% |
| Pig | ■ | 68.75% | 22.49% | 1.62% | 3.93% | 0.53% | 0.92% | 0.29% |

Supplementary Figure 3. The composition of four gene catalogs at Phylum level

**Supplementary Figure 4**

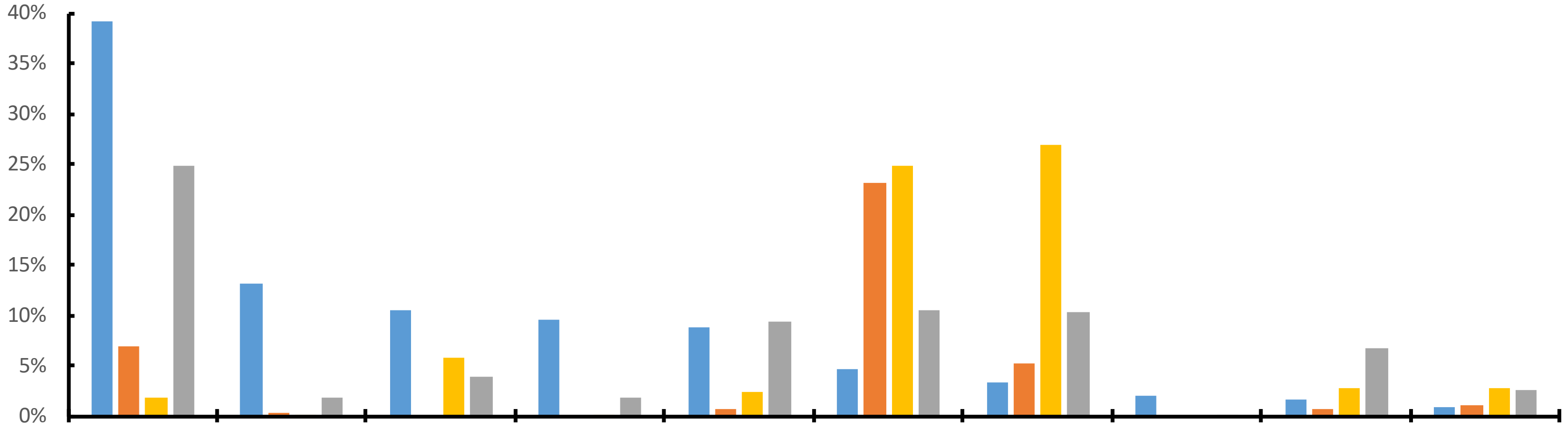

| | | Prevotella | Treponema | Butyrivibrio | Methanobrevib. | Ruminococcus | Bacteroides | Clostridium | Fibrobacter | Eubacterium | Roseburia |
|---|---|---|---|---|---|---|---|---|---|---|---|
| Rumen | ■ | 39.22% | 13.07% | 10.44% | 9.62% | 8.80% | 4.65% | 3.26% | 2.08% | 1.56% | 0.77% |
| Human | ■ | 6.90% | 0.28% | | | 0.72% | 23.20% | 5.26% | 0.00% | 0.59% | 0.98% |
| Mouse | ■ | 1.78% | 0.01% | 5.80% | 0.00% | 2.44% | 24.90% | 26.96% | 0.00% | 2.65% | 2.82% |
| Pig | ■ | 24.82% | 1.78% | 3.82% | 1.83% | 9.39% | 10.43% | 10.26% | 0.12% | 6.66% | 2.51% |

Supplementary Figure 4. The composition of four gene catalogs at genus level

**Supplementary Figure 5 “Mapped ratio on different ‘species’ catalog”**

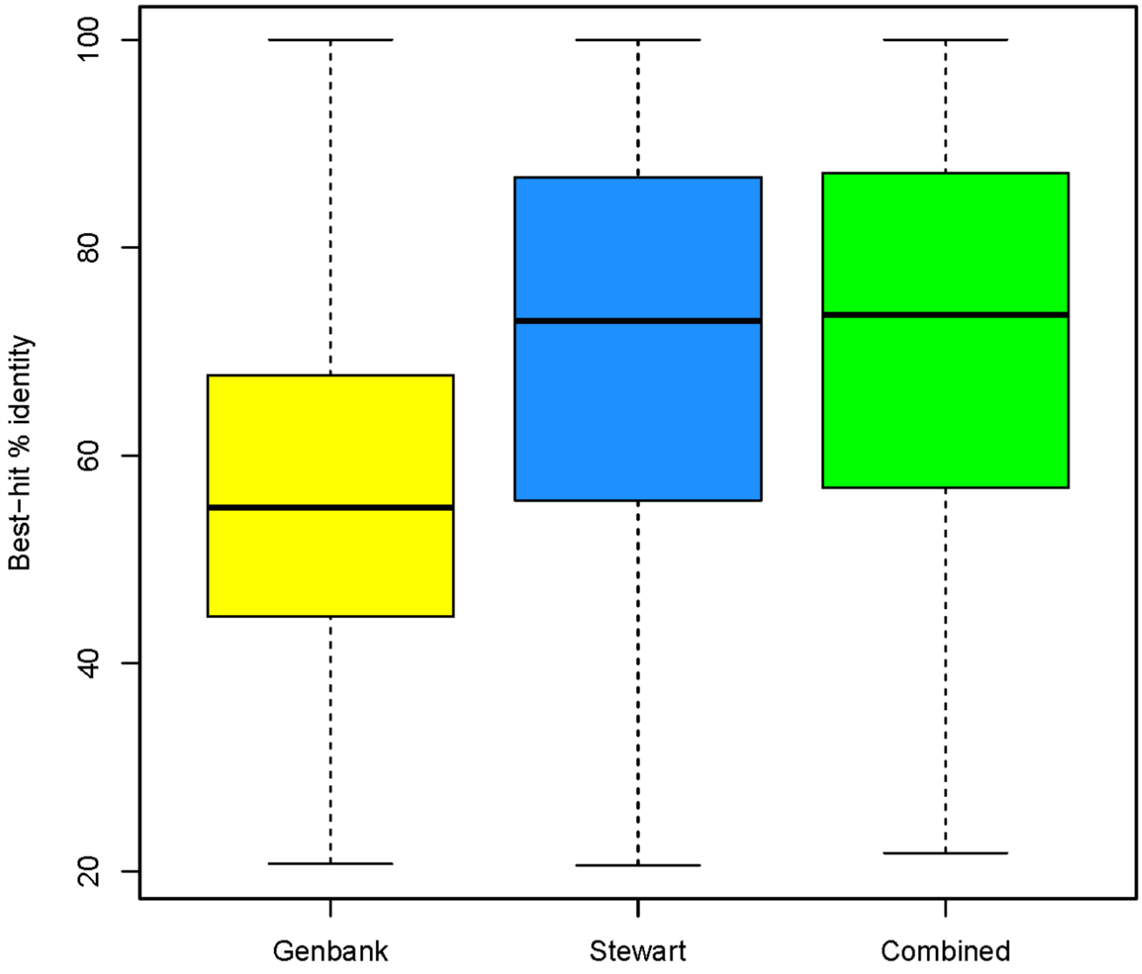

Supplementary Figure 5. CAZyme sequence diversity of our dataset against reference datasets. Distribution of identity percent obtained after best blast hits taking as queries the CAZymes in our catalog and searching for similarities in Genbank or Stewart et al. [1] datasets, alone or combined.

**Supplementary Figure 6**

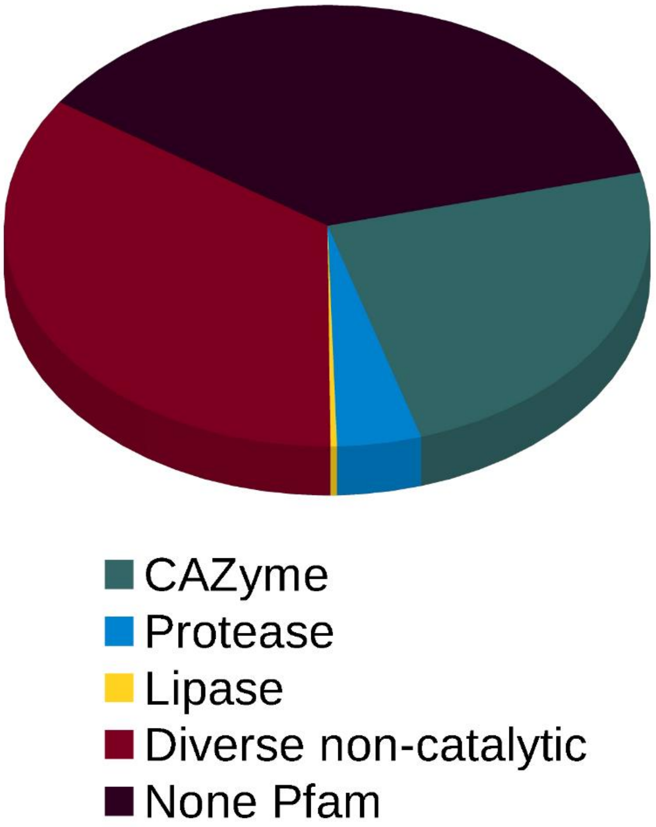

Supplementary Figure 6. Fonctional domains present in dockerin-containing proteins from the rumen gene catalog.

**Supplementary Figure 7**

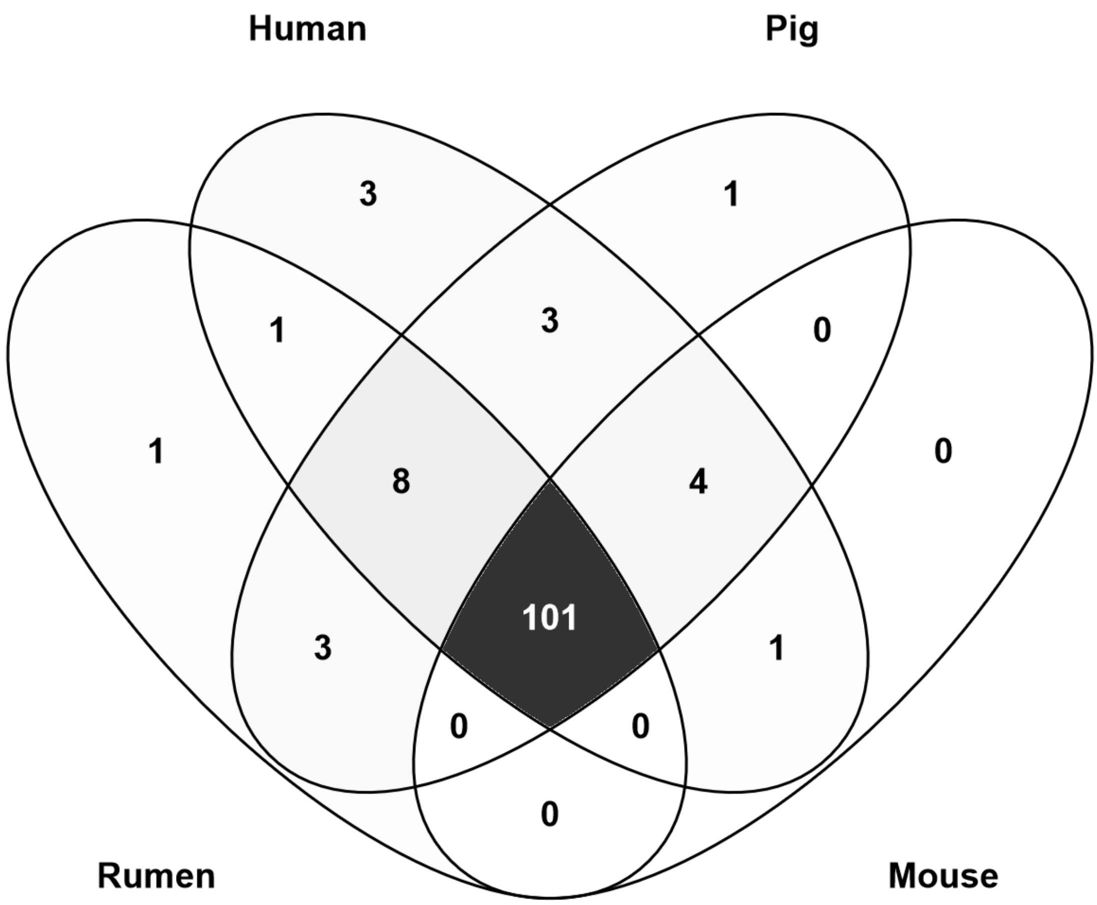

Supplementary Figure 7. Venn diagram of glycoside hydrolase families present in the gene catalogs from rumen, human, pig and mouse gut microbiota

**Supplementary Figure 8. HCA CAZy MAG**

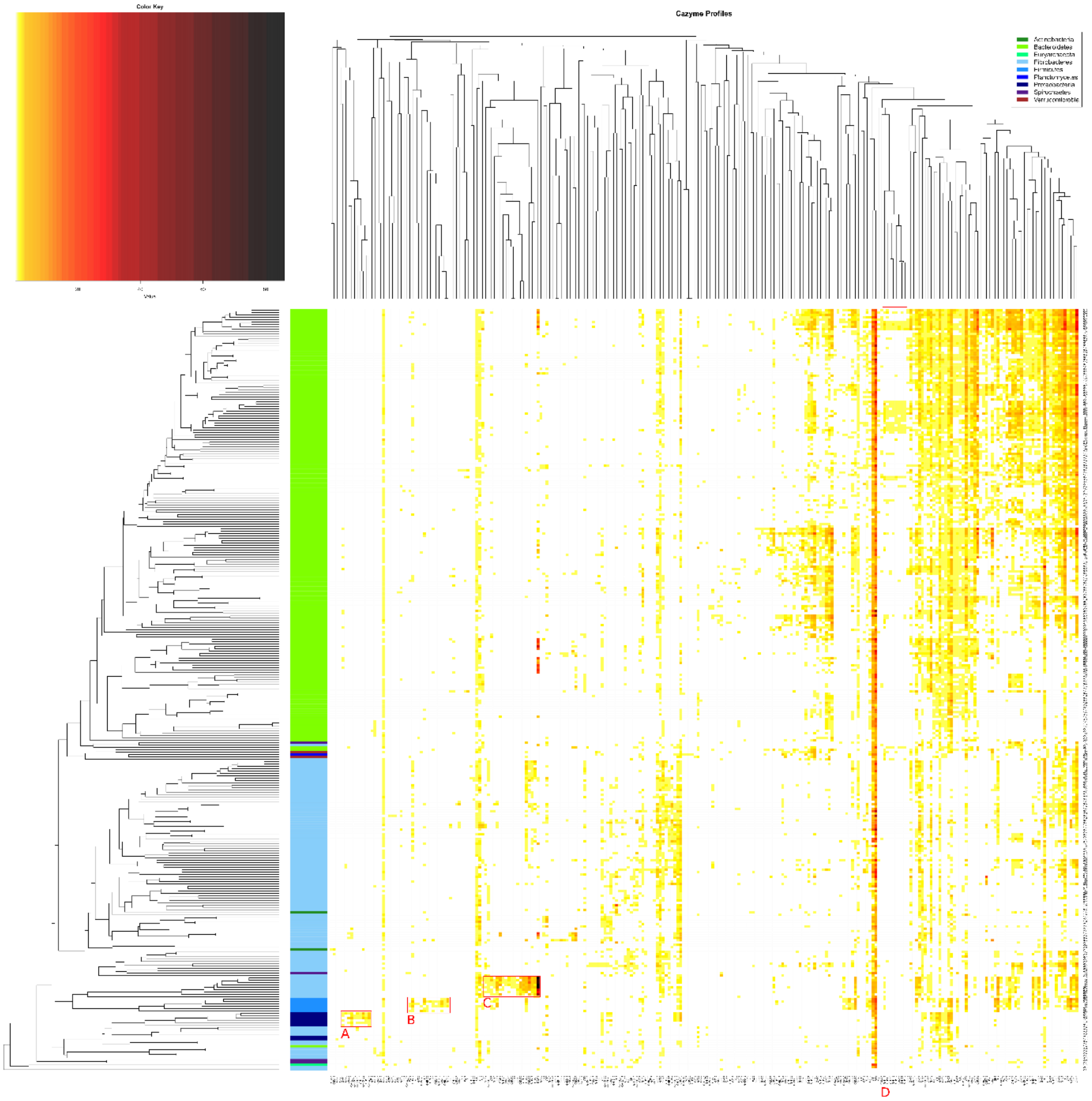

Supplementary Figure 8. Hierarchical clustering of 324 metagenomics species (MAG) according to their CAZyme profiles.

Rows display the 324 MAG with their identifiers on the right and predicted taxonomical phyla on the left (color-coded according to the top-right box). Columns display the abundance of glycoside hydrolase and polysaccharide lyase families (colored according to the top-left scale). Clustering was computed using the average-linkage method and Spearman's rank correlation. Four groups with specific CAZyme signatures are highlighted by red rectangles and labelled: (A) for *Proteobacteria* having specific GH13 subfamilies 19/32/37, and families GH84, GH103 and GH119; (B) for *Fibrobacteres* with many cellulases from families GH5, GH45 and GH55 and associated CBM11 and CBM30; (C) *Fimicutes* (likely from Ruminoccocus genus) with cellulosomal apparatus (dockerins and cohesins associated to various cellulases in famillies GH5, GH44, GH48 or GH124); and (D)

*Bacteroidetes* with recently discovered families GH137 to GH141 required to fully process the highly complex pectin component rhamnogalacturonan II

### Supplementary Figure 9. ‘similar to exp. PULs’

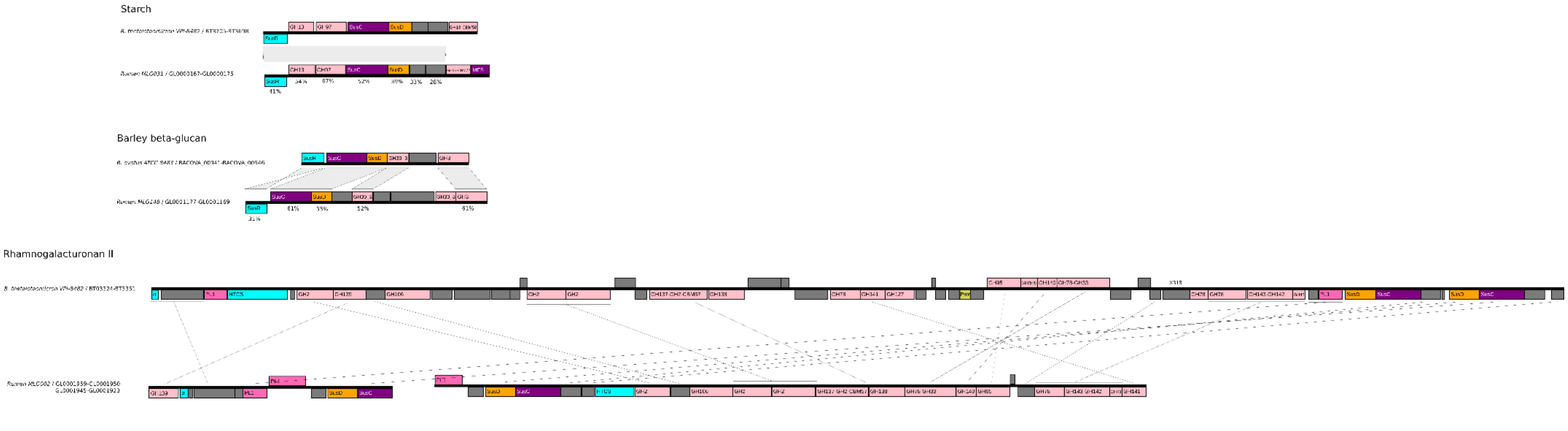

Supplementary Figure 9. Examples of Polysaccharide Utilization Loci (PULs) found bovine rumen metagenomic species (MAG) and their similarity to experimentally validated PULs with known target substrates. Rearrangements in PUL organization are indicated by grey shapes or segments linking gene homologs. Conservation between homologous proteins is illustrated by identity percentages obtained by BLASTP alignments.

**Common functions and influence of diet on the bovine rumen microbiome**

**Supplementary Figure 10 "NMDS gene counts by diet type (n= 77)"**

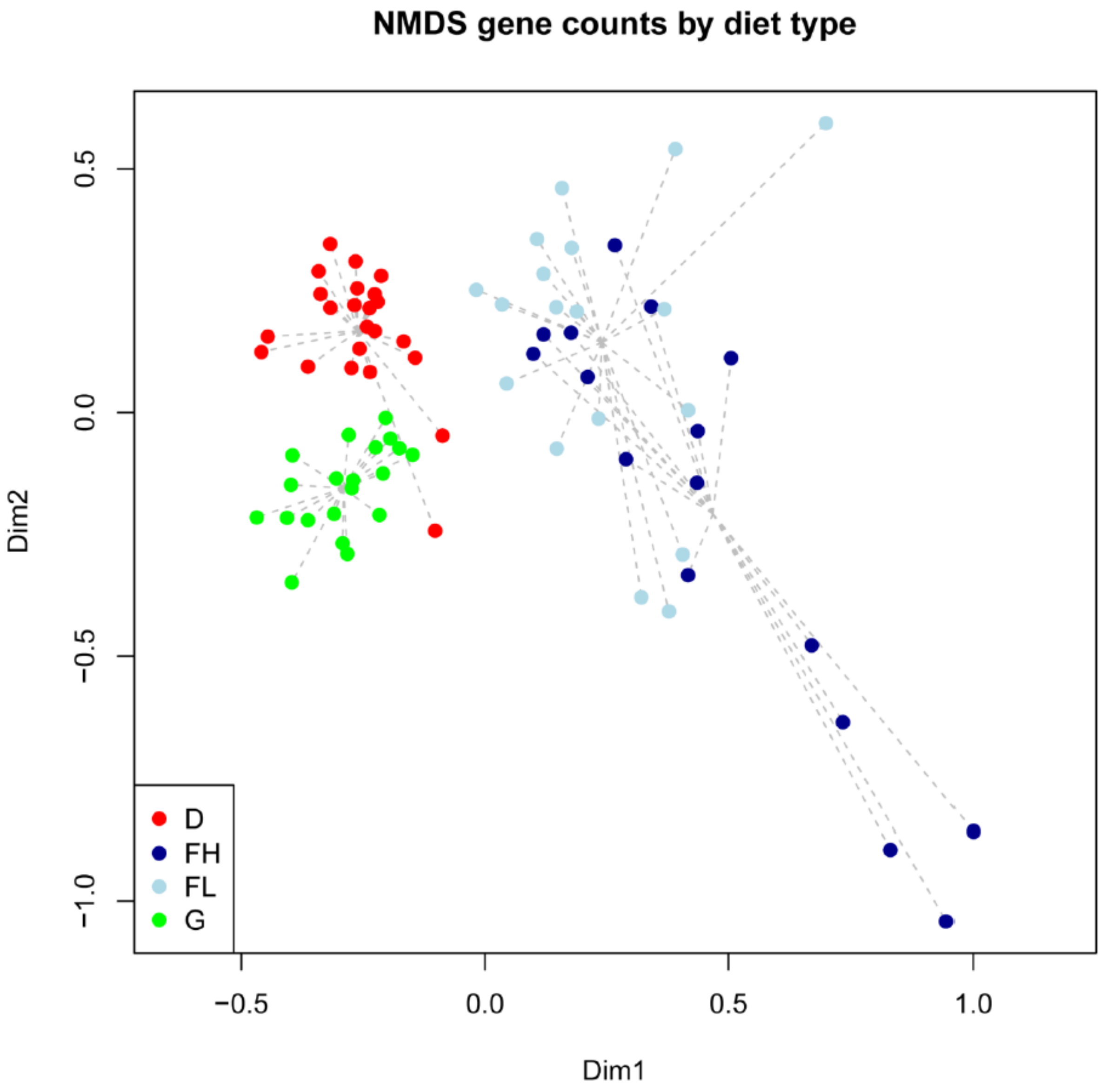

**Supplementary Figure 11. Overlapping number of Genus, CAZy and KO across diets (n= 77)**

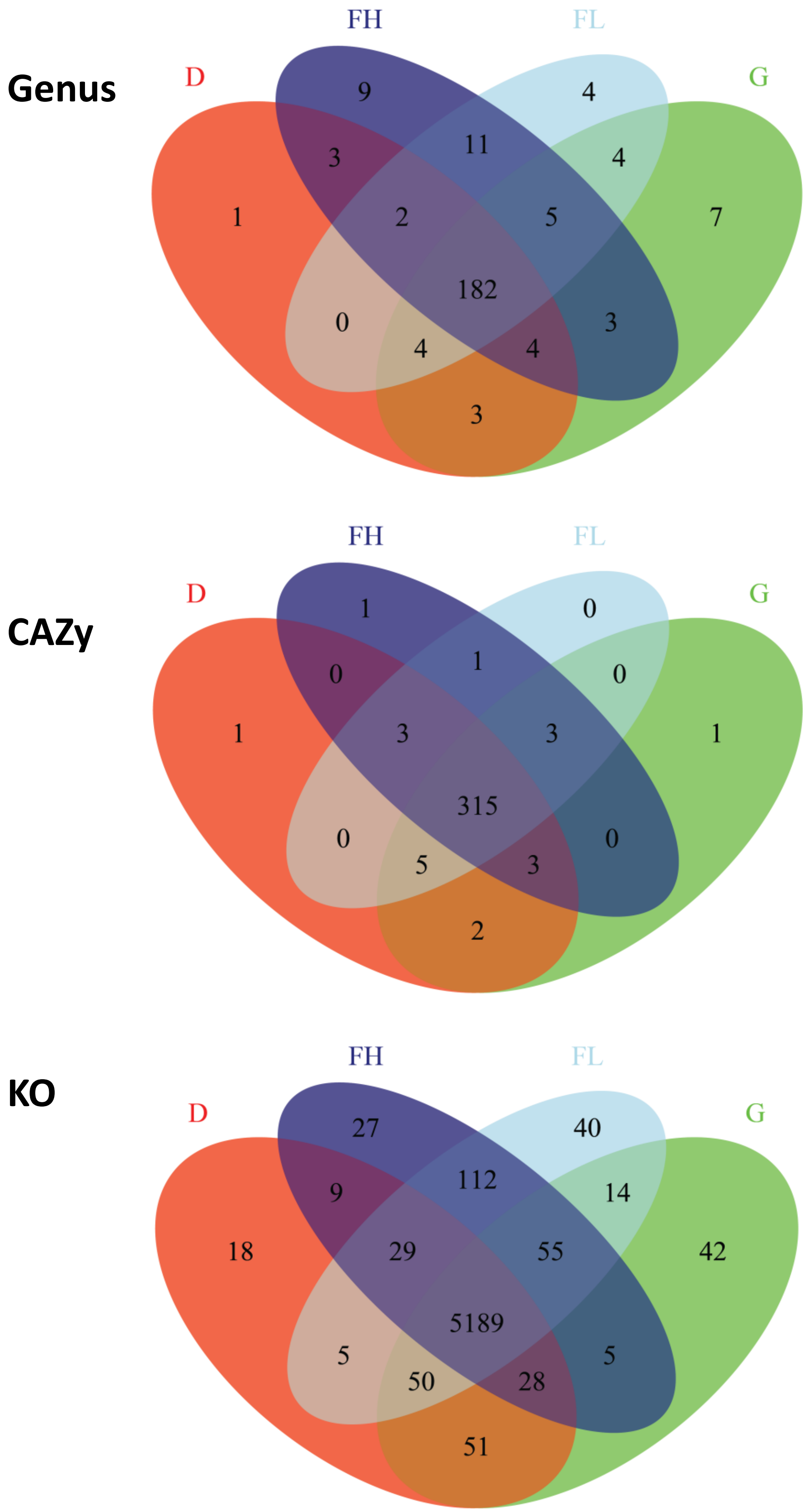

**Supplementary Figure 12**

a

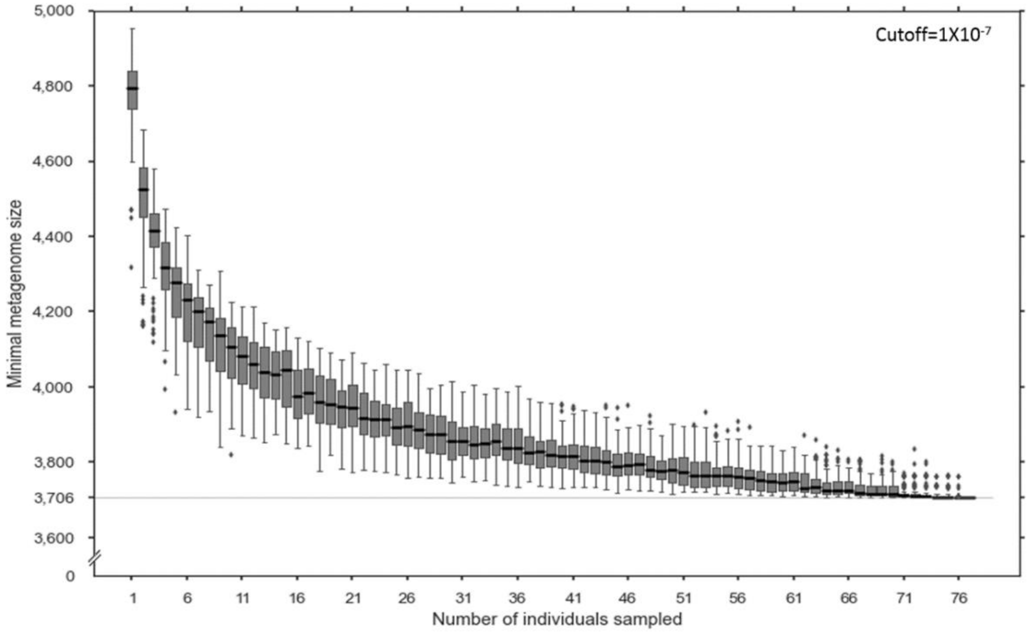

b

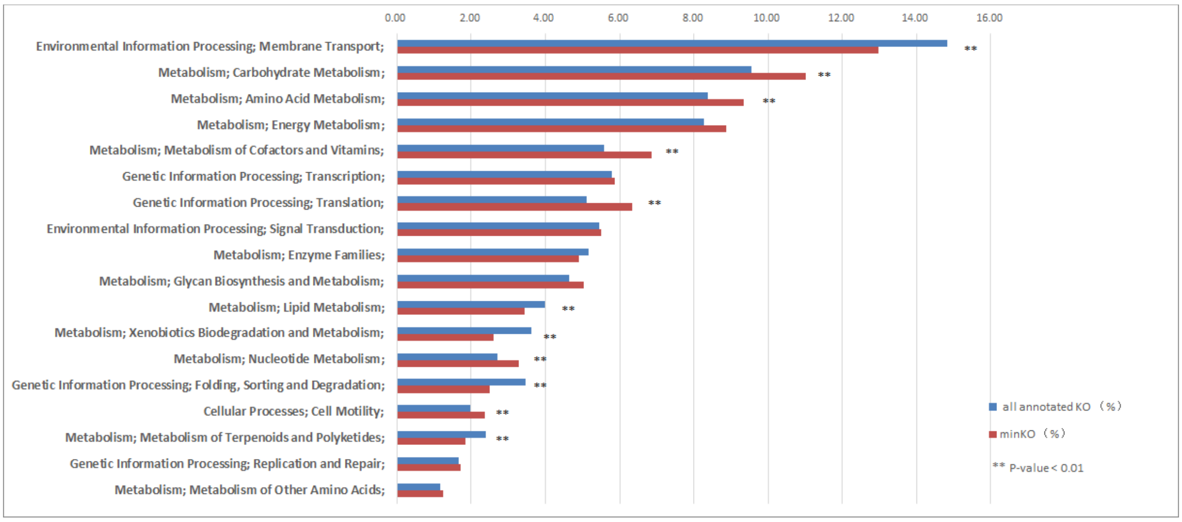

c

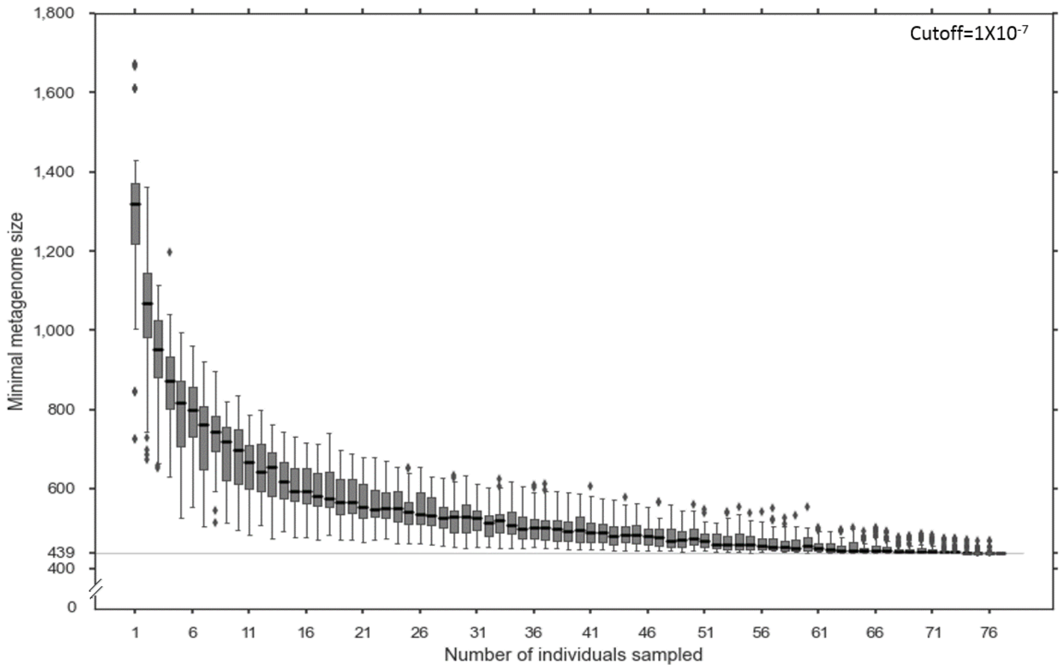

**d**

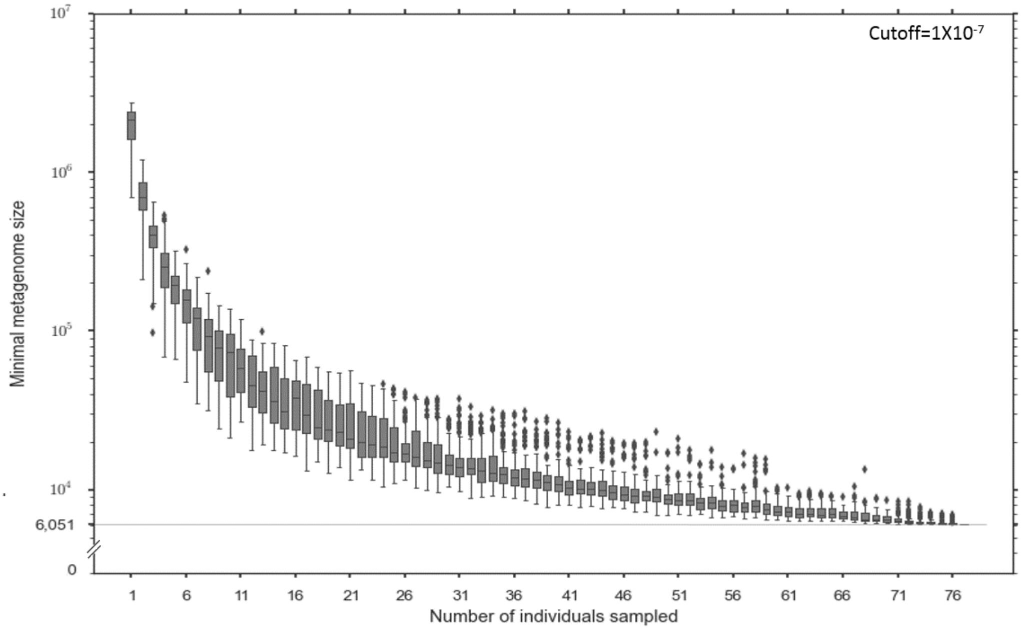

Supplementary Figure 12. Size of the shared microbiome features among cattle fed four different diets using different calculation methods than that of Figure 2 for comparison (see Methods section).
(a) The size of the minimal metagenome at KO levels (cutoff= $1X10^{-7}$). There are 5893 functionally annotated KO levels and a minimal set of 3706 functions was found for the 77 individuals sampled. (b) Change in the rate of all annotated KO and minimal KO in the second function level. (c) The size of the minimal metagenome at CAZy levels (cutoff= $1X10^{-7}$). There are 1974 functionally annotated CAZy levels and the minimal set found was of 439 functions. (d) The size of minimal metagenome at gene levels (cutoff= $1X10^{-7}$), 6051 genes was the minimal set for the 77 individuals sampled.

**Supplementary Figure 13 “cluster distrib. figures”**

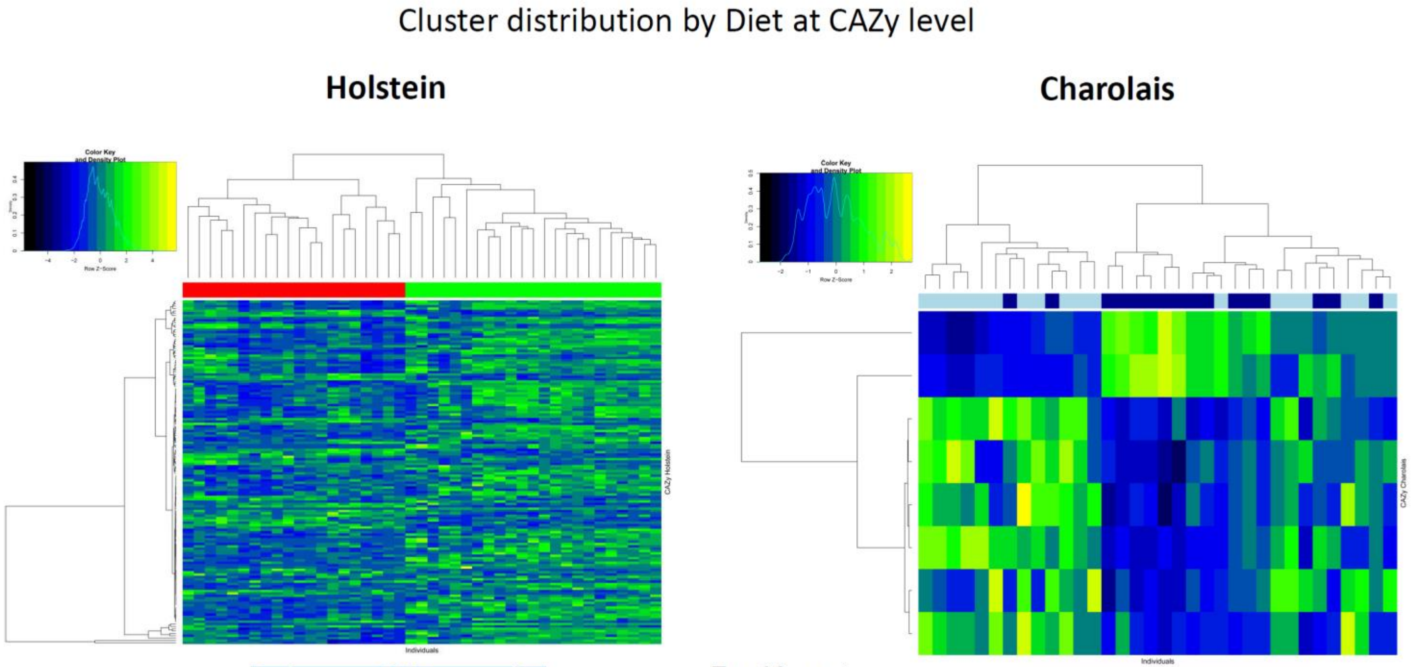

**Top 10 most abundant DA CAZy**

| G | CAZy | D |
|---|---|---|
| | GT2 | ↑ |
| | GT4 | ↑ |
| | CE1 | ↑ |
| | CBM6 | ↑ |
| | DOC1 | ↑ |
| | GH13 | ↑ |
| | CBM50 | ↑ |
| | GH9 | ↑ |
| | GH10 | ↑ |
| | GT35 | ↑ |

| FL | CAZy | FH |
|---|---|---|
| | GH73 | ↑ |
| | GH24 | ↑ |
| ↑ | GH65 | |
| ↑ | GT66 | |
| ↑ | GH113 | |
| ↑ | PL9_2 | |
| ↑ | CBM66 | |
| ↑ | GH13_18 | |

Supplementary Figure 13 (A) Cluster distribution in Holstein and Charolais breeds based on CAZy families. For Holstein red label indicates dairy cow (D, n= 23) diet and green indicates grazing (G, n= 20) diet. For Charolais, dark blue indicates feedlot high starch + linseed (FH, n= 16) diet and light blue indicates feedlot low starch (FL, n=18) diet. Changes in differentially abundant families for each breed are shown in the tables below the graph (arrows indicate higher abundance). The complete list is available in Supplementary Table 9.

#### Cluster distribution by Diet at KO level

**Holstein** **Charolais**

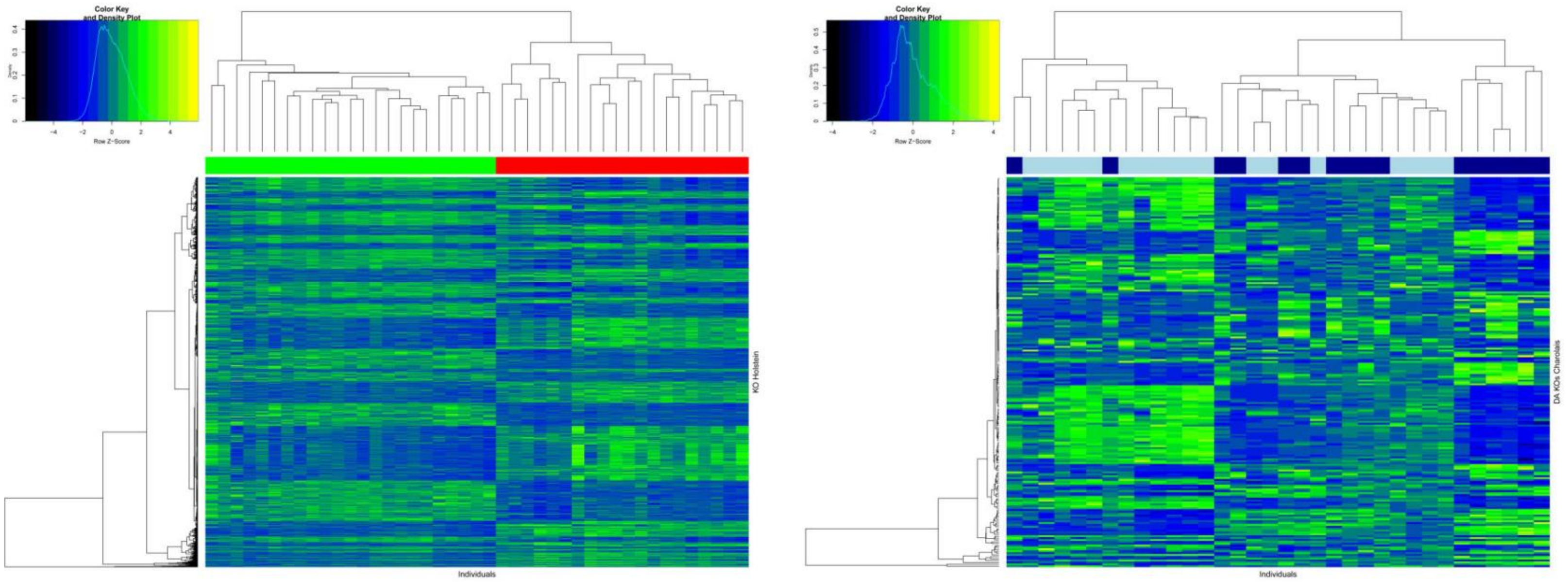

**10 Top DA KO**

| Holstein | | Charolais | |
|---|---|---|---|
| K00754 | K02003 | K00680 | K02982 |
| K02004 | K01179 | K11527 | K03183 |
| K00599 | K03657 | K00721 | K12063 |
| K00936 | K01190 | K07485 | K02926 |
| K01897 | K06180 | K01652 | K03496 |

Supplementary Figure 13 (B) Cluster distribution in Holstein and Charolais breeds based on KEGG orthology. For Holstein red label indicates dairy cow (D) diet and green indicates grazing (G) diet. For Charolais, dark blue indicates feedlot high starch + linseed (FH) diet and light blue indicates feedlot low starch (FL) diet. Changes in the 10 most differentially abundant KO for each breed are shown below the graph. The complete list is available in Supplementary Table 16.

#### Cluster distribution by Diet at Genus level

**Holstein**

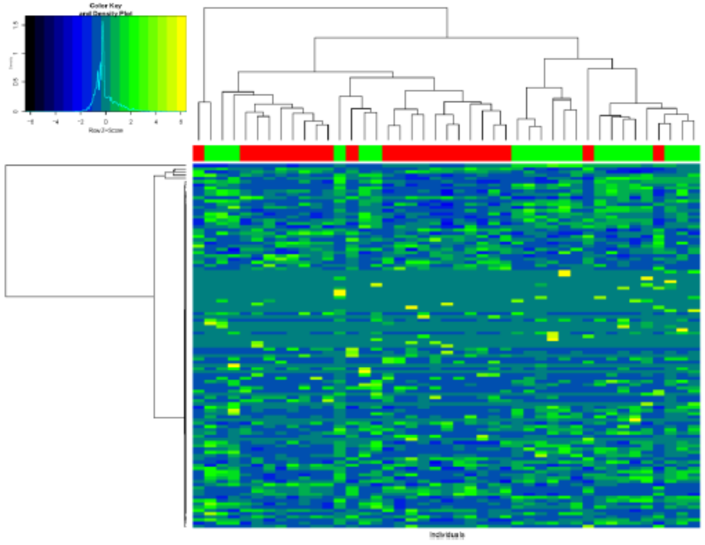

**Charolais**

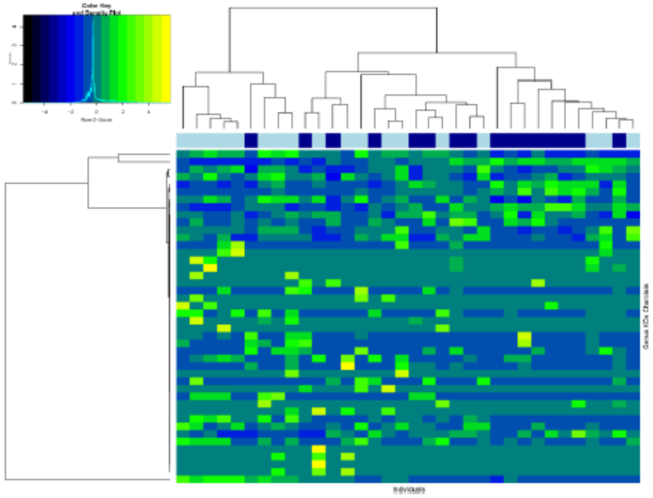

| D | Genus | G |
|---|---|---|
| | *Prevotella* | ↑ |
| | *Butyrivibrio* | ↑ |
| ↑ | *Fibrobacter* | |
| | *Methanobrevibacter* | ↑ |
| ↑ | *Ruminococcus* | |
| ↑ | *Bifidobacterium* | |
| ↑ | *Roseburia* | |
| | *Selenomonas* | ↑ |
| ↑ | *Acidaminococcus* | |
| | *Methanosphaera* | ↑ |
| ↑ | *Lactobacillus* | |
| | *Desulfovibrio* | ↑ |

**Top most abundant DA Genus**

| FH | Genus | FL |
|---|---|---|
| | *Bacteroides* | ↑ |
| | *Fibrobacter* | ↑ |
| ↑ | *Butyrivibrio* | |
| | *Methanosphaera* | ↑ |
| | *Streptococcus* | ↑ |
| ↑ | *Acidaminococcus* | |
| ↑ | *Lactobacillus* | |
| | *Faecalibacterium* | ↑ |
| ↑ | *Dehalococcoides* | |
| ↑ | *Haemophilus* | |

Supplementary Figure 13 (C) Cluster distribution in Holstein and Charolais breeds based on microbial genera. For Holstein red label indicates dairy cow (D) diet and green indicates grazing (G) diet. For Charolais, dark blue indicates feedlot high starch + linseed (FH) diet and light blue indicates feedlot low starch (FL) diet. Changes in differentially abundant genera are shown in the tables below the graph (arrows indicate higher abundance). The complete list is available in Supplementary Table 10.

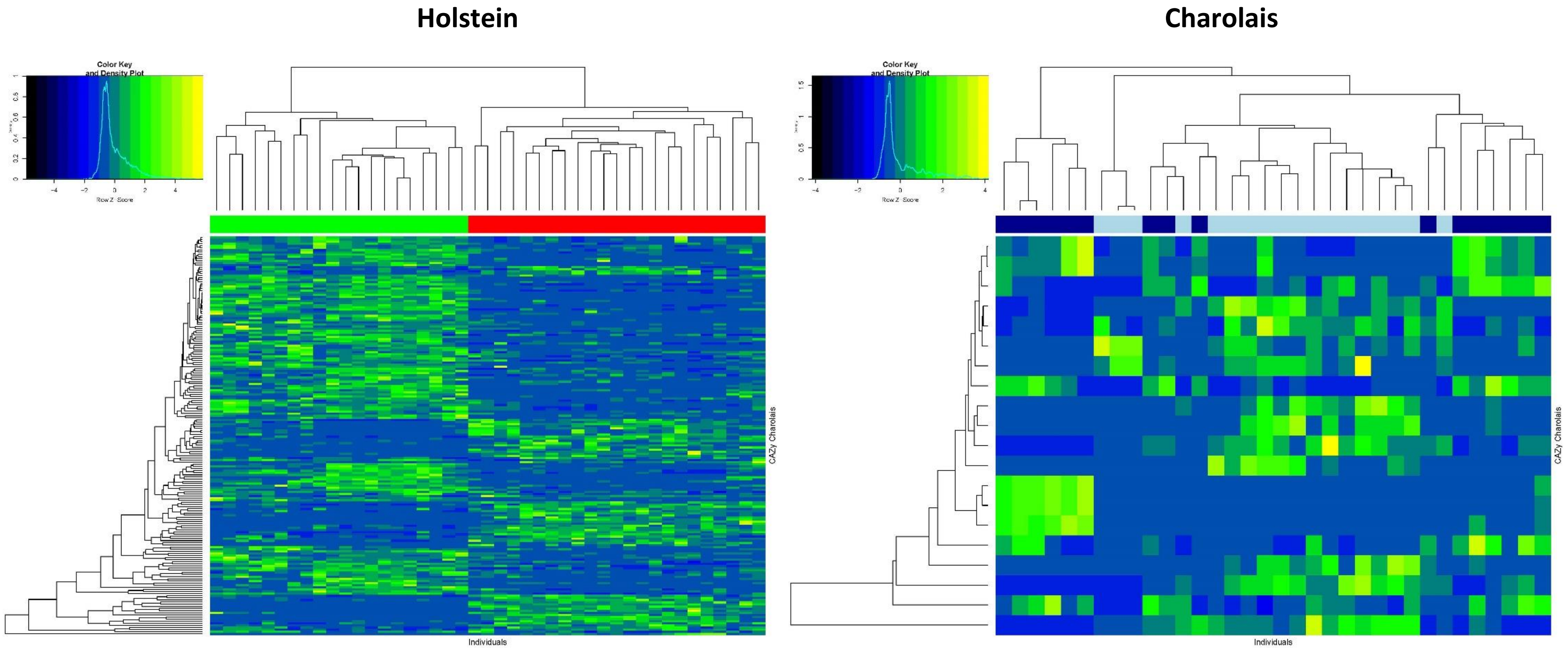

Supplementary Figure 13 (D) Cluster distribution in Holstein and Charolais breeds based on MAGs. For Holstein, red label indicates dairy cow (D) diet and green indicates grazing (G) diet. For Charolais, dark blue indicates feedlot high starch + linseed (FH) diet and light blue indicates feedlot low starch (FL) diet. The complete list is available in Supplementary Table 10.

**Supplementary Figure 14**

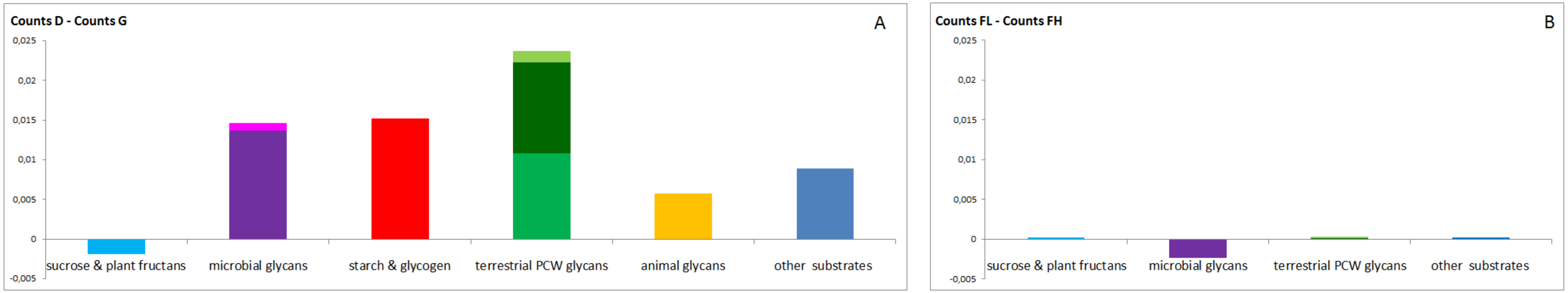

**Supplementary Figure 14.** Effect of diet on the abundance of CAZy families in the bovine rumen metagenome. Difference of abundance of CAZy families, clustered by substrate categories, between the D (n= 23) and G (n= 20) samples (A), and between the FL (n= 16) and FH (n= 18) samples (B). Only the most significant differentially abundant catabolic CAZy families (glycoside hydrolases, polysaccharide lyases, and family 35 of glycosyl transferases, of which the members are able to cleave osidic linkages of starch and glycogen by phosphorolysis) and their associated non-catalytic modules were taken into account (adjusted pvalue < 0.05). Color legend: pale purple, fungal glycans; dark purple, bacterial glycans; pale green, pectin; medium green, cellulose; dark green, hemicellulose. Polyspecific families containing members acting on various types of substrates are assigned to 'other substrates'

#### Antibiotic resistance genes

##### Supplementary Figure 15. ARG

a)

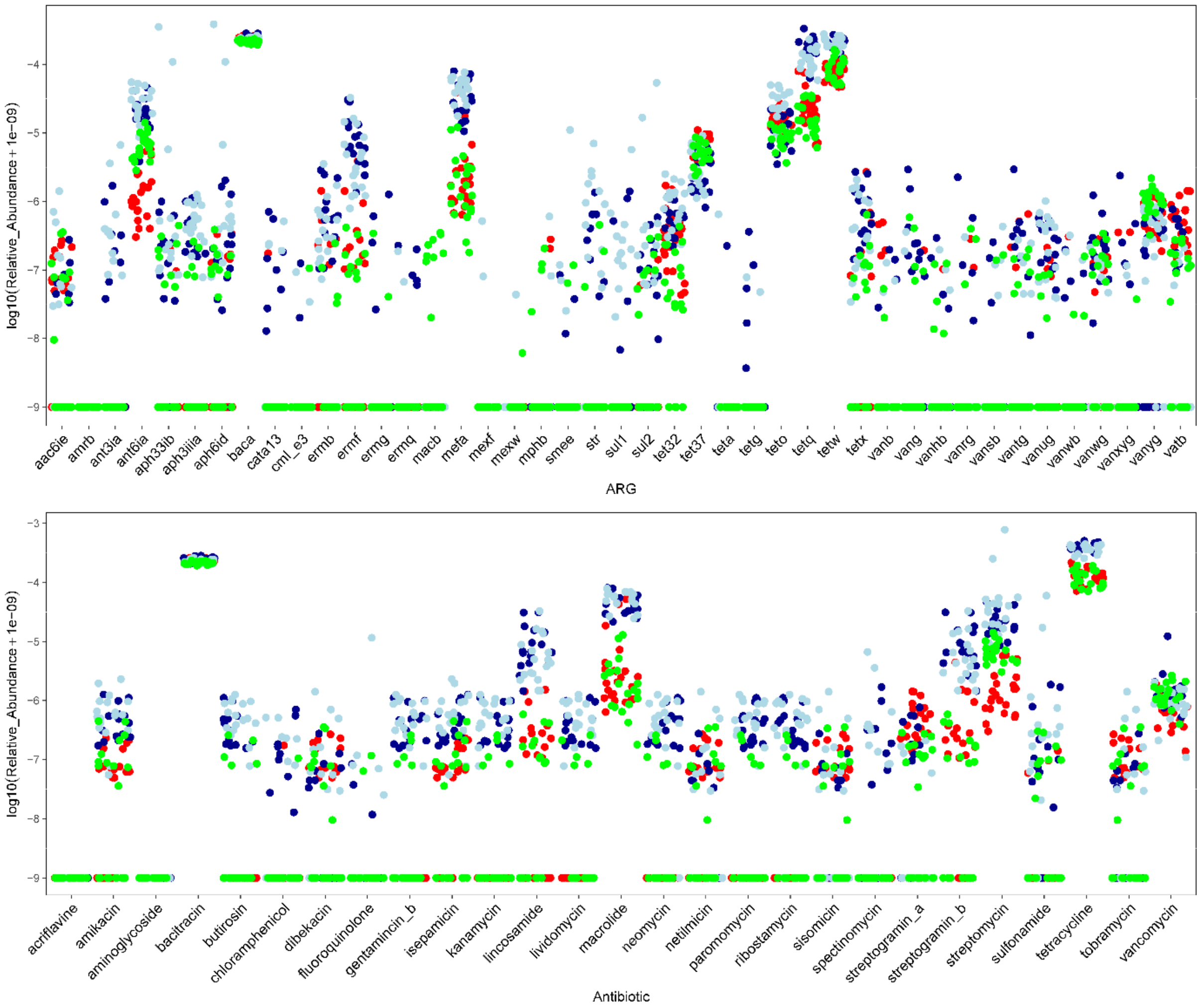

b)

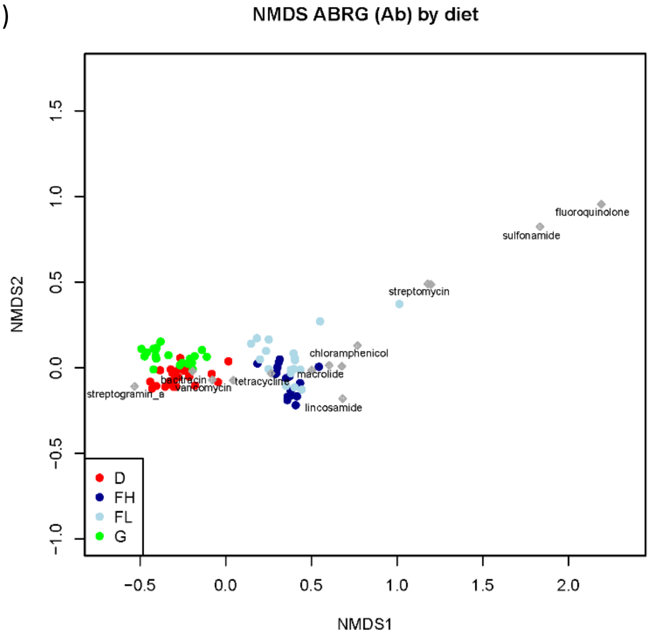

Supplementary Figure 15. Prevalence of antibiotic resistance genes (ARGs) in the 77 samples cohort. a) Relative abundances (log10 scale) of ARGs (top) and antibiotic types (bottom) found in each individual. In every column on the x axis each dot represents an animal, with colors according to diets: grazing (G; green), dairy (D; red), fattening high-start (FH; dark blue) and fattening low-starch (FL; light blue). The higher the vertical position of the dots on the y axis, the higher the relative abundance of the ARGs.
b) NMDS biplot of the ARGs showing a separation between beef (FH and FL diets) and dairy breeds (D and G diets). Grey diamonds represent the individual ARG ordination onto the two-dimensional space, with the names of a subset of ARGs given in black.

### Methods

#### Construction of metagenomic species (MAG) and taxonomic assignment

**Supplementary Figure 16**

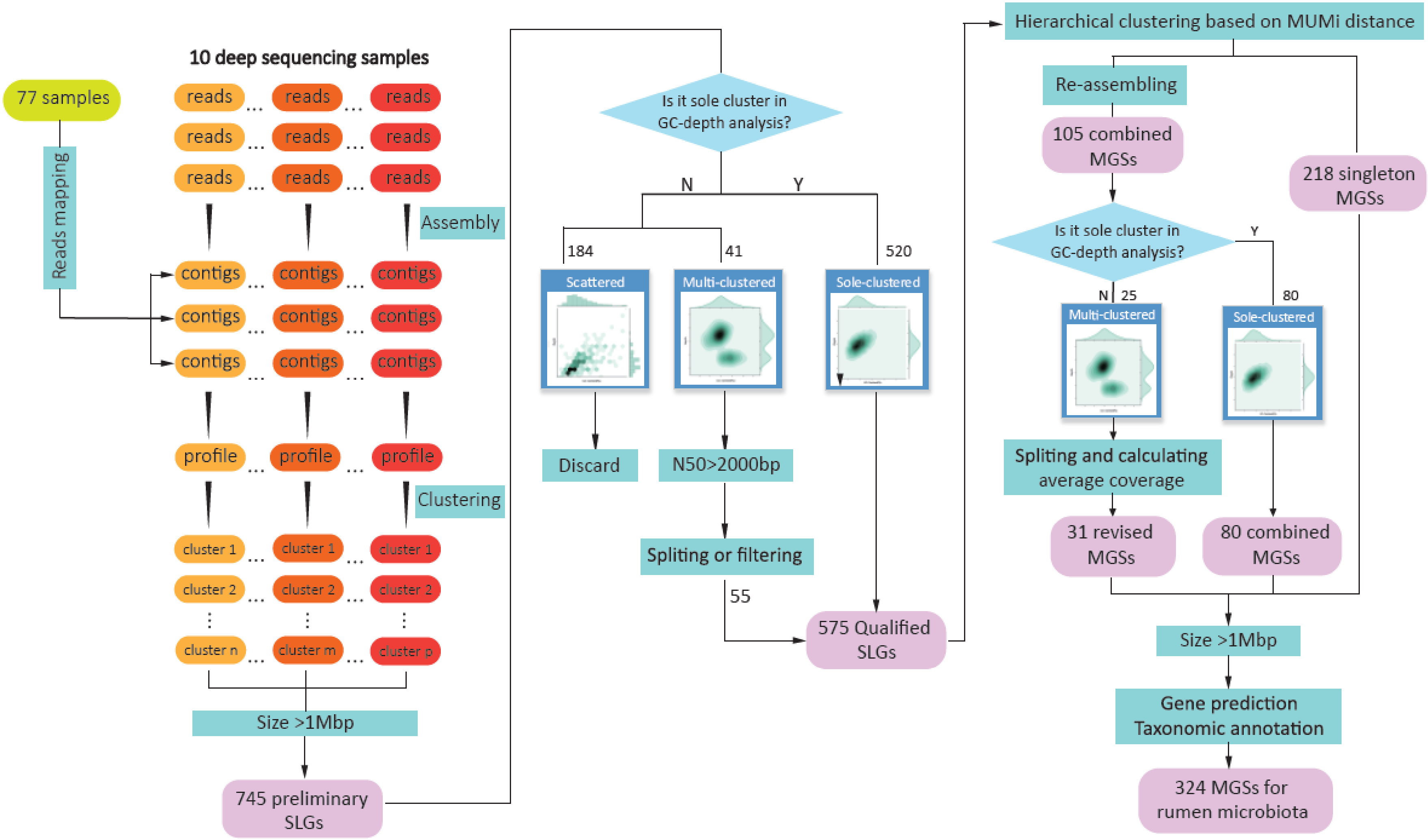

Supplementary Figure 16. Diagram describing the approach used for the construction of rumen metagenomics clusters

**Supplementary Figure 17**

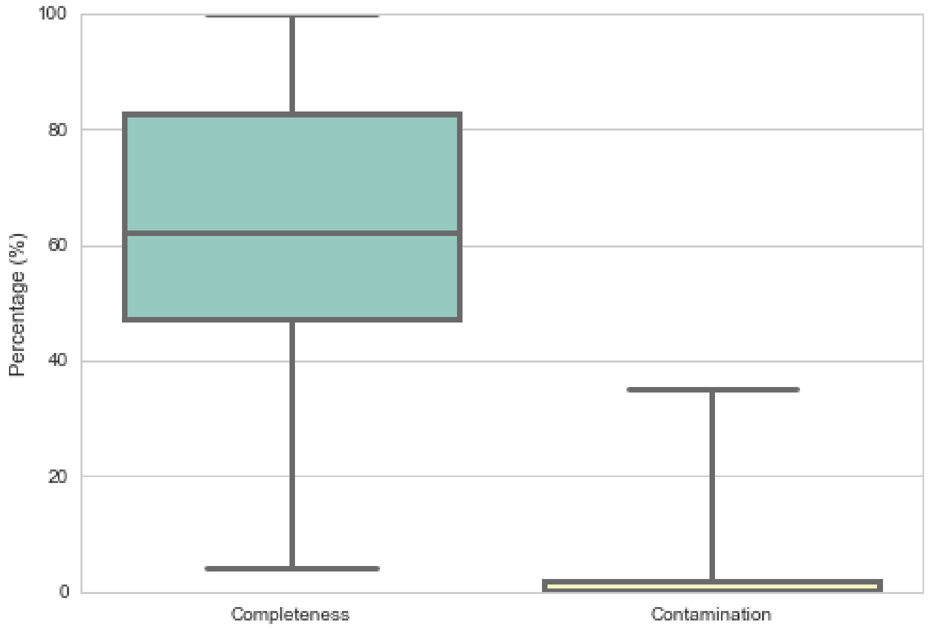

**Supplementary Figure 17. Percentage of 324 MAGs completeness and contamination.**
The median percentage of completeness and contamination reached 62.48% and 2.62%, respectively (see Supplementary Note 1) showing that the 324 MAGs have high completeness and low contamination.

**Supplementary Figure 18**

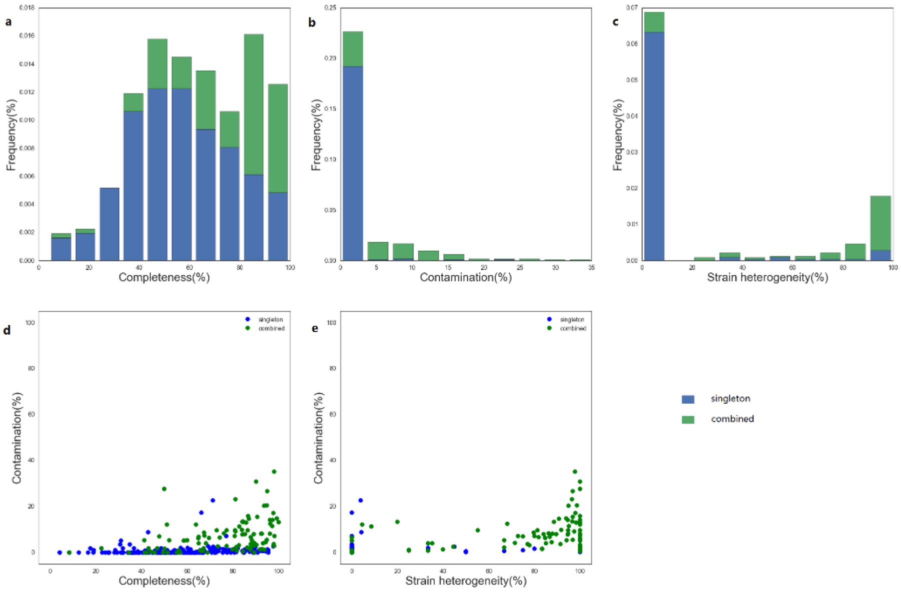

Supplementary Figure 18. Completeness, Contamination and Strain heterogeneity in singleton (blue) and combined (red) MAGs.

(a)Proportion of completeness between combined and singleton MAGs. Most MAGs show a completeness greater than 40%. (b) Frequency of contamination between combined and singleton MAGs. Contamination in most MAG is 0, the rest of MAGs contamination is mainly due to combined MAGs. (c) proportion of strain heterogeneity between combined and singleton MAGs. Strain heterogeneity in most singleton MAG is 0, whereas most combined MAGs have a strain heterogeneity greater than 70%. (d) When MAG completeness is greater than 40% then most of MAGs contamination is less than 10%. (e) When the MAGs contamination greater than 10%, the contamination of MAGs (mostly combined MAG) basically come from strain heterogeneity.

#### Quality assessment and taxonomic annotation of MAGs

**Supplementary Figure 19**

a)

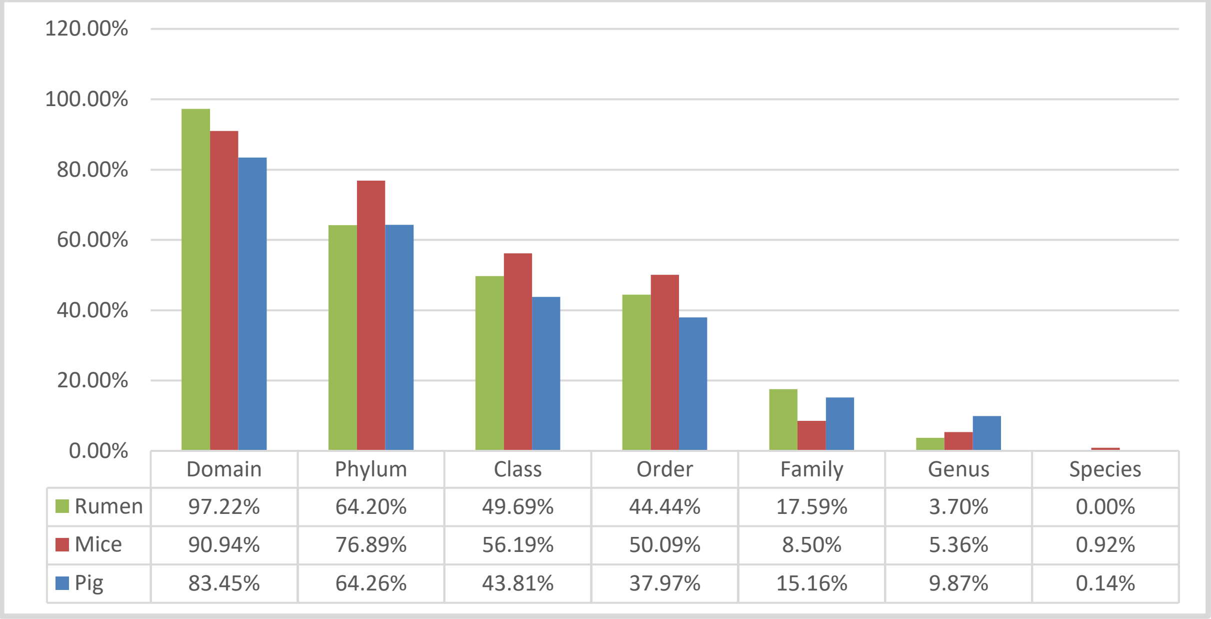

| | Domain | Phylum | Class | Order | Family | Genus | Species |
|---|---|---|---|---|---|---|---|
| Rumen | 97.22% | 64.20% | 49.69% | 44.44% | 17.59% | 3.70% | 0.00% |
| Mice | 90.94% | 76.89% | 56.19% | 50.09% | 8.50% | 5.36% | 0.92% |
| Pig | 83.45% | 64.26% | 43.81% | 37.97% | 15.16% | 9.87% | 0.14% |

b)

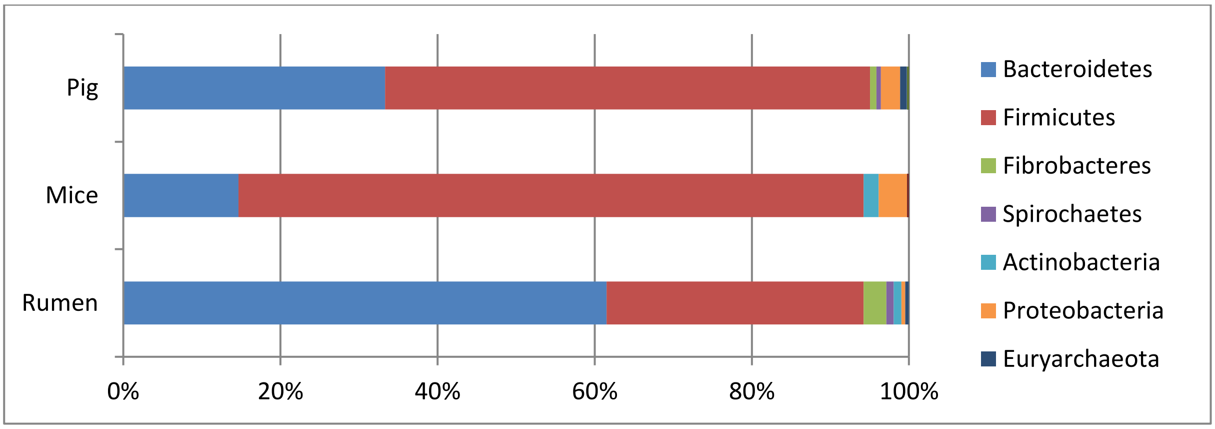

c)

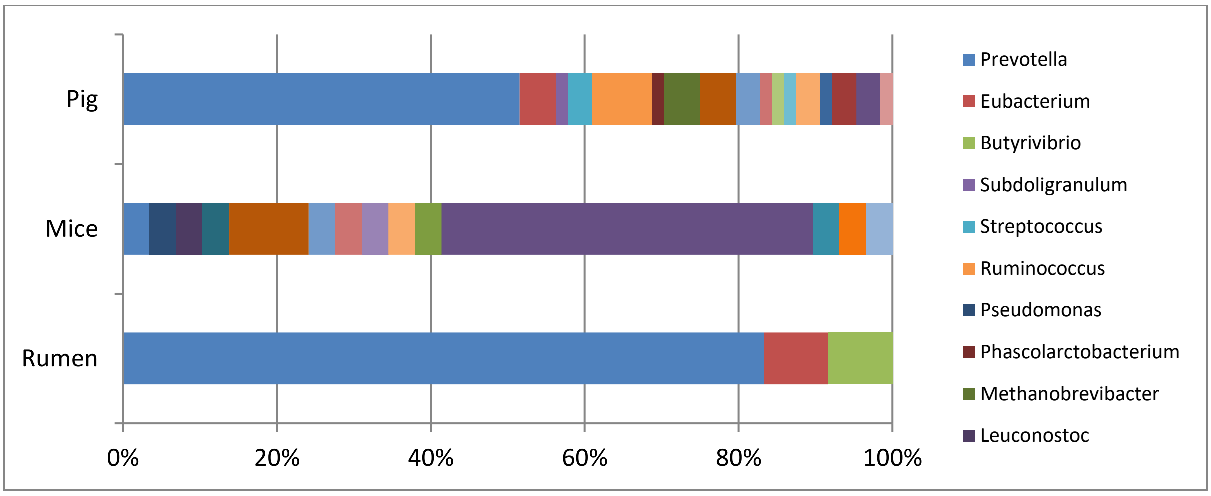

Supplementary Figure 19. The distribution of MAG from the bovine rumen and pig and mice feces metagenomes at different taxonomic levels (a). MAG distribution at phylum (b) and genus (c) level.

**Supplementary Figure 20. Relationship between coverage and depth for MAG**

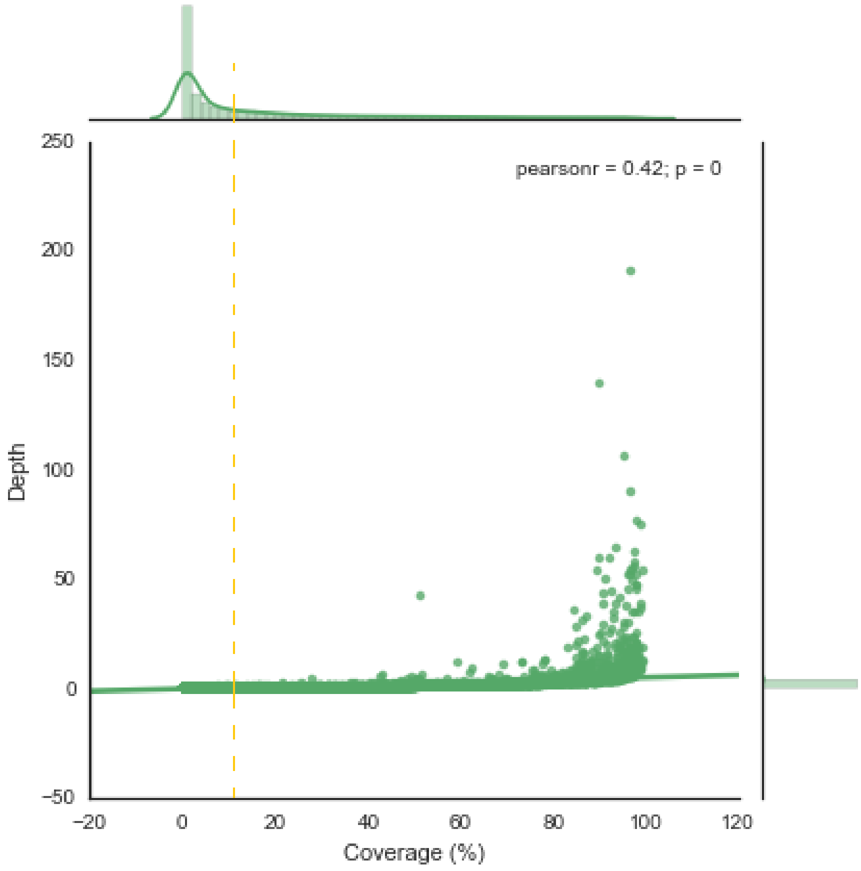

Supplementary Figure 20. Relationship between coverage and depth for MAG.  Each point represents one MAG in one of the 77 samples. Histogram in the top and right-hand indicates the fraction of coverage and depth, respectively. When the coverage is less than 10%, then the value of depth is close to 0.

**Supplementary Table 1**

Supplementary Table 1. Summary information of the bovine rumen prokaryotic gene catalog described in this study

| Total number of genes | Number of complete genes | Total length (bp) | Average length (bp) | N50 (bp) | N90 (bp) | Max length | Min length | Taxonomic Annotation (Phylum Level) | Taxonomic Annotation (Genus Level) | KEGG Annotation | Cazy Annotation |
|---|---|---|---|---|---|---|---|---|---|---|---|
| 13,825,880 | 5,410,738 | 9,896,715,705 | 715.81 | 906 | 396 | 40,812 | 102 | 5,899,724 (42.67%) | 1,205,758 (8.72%) | 5,947,496 (43.02%) | 174,226 (3.22%) |

**Supplementary Table 2**

Supplementary Table 2. Summary information of the bovine rumen gene catalog described by Hess et al. (2011) pre- and post-filtering

| | Number of genes | Number of bases | Average length | Max length | Min length | N50 (bp) | N90 (bp) |
|---|---|---|---|---|---|---|---|
| Published data | 2,547,270 | 1,379,576,700 | 541.59 | 319,74 | 60 | 744 | 261 |
| Filtered data | 2,461,518 | 1,218,405,383 | 494.98 | 319,74 | 100 | 681 | 231 |

**Supplementary Table 3. "Supplementary Table 3 - Excel file"**

**Supplementary Table 3**. Similarity between MAGs from this work, Hungate1000 genomes and MAGs from Stewart et al. [1]

**Supplementary Table 4. "Supplementary Table 4 - Excel file"**

**Supplementary Table 4.** Metagenomic data production from bovine rumen samples

**Supplementary Table 5. "Supplementary Table 5 -Excel file"**

**Supplementary Table 5.** Number of genes and taxonomic annotation information for bovine rumen and human, pig and mouse feces gene catalogs

**Supplementary Table 6**

Table 6. Detailed annotation information at Genus level for rumen, pig and mice catalogs

| Genus | Rumen | Pig | Mice |
|---|---|---|---|
| *Prevotella* | 39.22% | 24.85% | 1.79% |
| *Treponema* | 13.07% | 1.78% | 0.01% |
| *Butyrivibrio* | 10.44% | 3.82% | 5.83% |
| *Methanobrevibacter* | 9.62% | 1.83% | 0.00% |
| *Ruminococcus* | 8.80% | 9.40% | 2.45% |
| *Bacteroides* | 4.65% | 10.44% | 25.01% |
| *Clostridium* | 3.26% | 10.27% | 27.08% |
| *Fibrobacter* | 2.08% | 0.12% | 0.00% |
| *Eubacterium* | 1.56% | 6.67% | 2.66% |
| *Roseburia* | 0.77% | 2.51% | 2.83% |

**Supplementary Table 7. Excel file "Supplementary Table 7_CAZy-counts_human_rumen_mouse_pig_catalogs"**

**Supplementary Table 7.** Proportion of carbohydrate active enzymes (CAZy) in the rumen microbiome of cattle and gut microbiomes of human, pig and mouse.

**Supplementary Table 8. Excel file "Supplementary Table 8_DA_Fibrobacter_CAZy"**

**Supplementary Table 8.** Effect of diet on the abundance of CAZy genes assigned to *Fibrobacter succinogenes* in the bovine rumen metagenome. Gene counts of the differentially abundant genes in the D and G samples (spreadsheet 'Holstein'), and in the FL and FH samples (spreadsheet 'Charolais'). Similarity of 1262 CAZy Fibrobacteres genes enumerated in the catalogue to *Fibrobacter succinogenes* S85.

**Supplementary Table 9. Differential abundance analysis**

Supplementary Table 9. Number of differentially abundant Genus, MAG, CAZy and KO within each breed detected by differential abundant analysis.

| Breed | Genus | MAG | CAZy | KO |
|---|---|---|---|---|
| Holstein | 106 | 188 | 146 | 2445 |
| Charolais | 44 | 20 | 8 | 208 |

**Supplementary Table 10. Excel file "Supplementary Table 10" significant differences CAZy**

**Supplementary Table 10.** Effect of diet on the abundance of CAZy families in the bovine rumen metagenome. Gene counts of the differentially abundant families of glycoside hydrolases, polysaccharide lyases, family 35 of glycosyl transferases, carbohydrate-binding modules, dockerins and cohesins in the D and G samples (sheet 'Holstein'), and in the FL and FH samples (sheet 'Charolais'). The targeted substrates are indicated for each family.

**Supplementary Table 11. Excel file "Supplementary Table 11_DA_Genus_MAG"**

**Supplementary Table 11.** Effect of diet on the abundance of genera and MAGs in the bovine rumen metagenome.

**Supplementary Table 12. "Vector fitting Table"**

#### Holstein (only significant results are presented)

| Pheno | Dim1 | Dim2 | r2 | Pr(>r) |
|---|---|---|---|---|
| LW_kg | -0.41358 | -0.91047 | 0.3533 | 0.0004 |
| total_VFA | -0.27517 | 0.96140 | 0.2799 | 0.0030 |
| C2 | -0.31894 | 0.94778 | 0.2711 | 0.0038 |
| C3 | 0.01138 | 0.99994 | 0.2909 | 0.0025 |
| iC4 | 0.99504 | -0.09952 | 0.3326 | 0.0009 |
| C4 | -0.57452 | 0.81849 | 0.3797 | 0.0001 |
| iC5 | 0.94064 | -0.33941 | 0.1487 | 0.0541 |
| C5 | 0.26250 | 0.96493 | 0.1321 | 0.0787 |
| C6 | -0.97602 | -0.21767 | 0.3948 | 0.0002 |
| C2_C3 | -0.47642 | -0.87922 | 0.2946 | 0.0019 |
| total_protozoa | -0.97810 | -0.20814 | 0.5500 | 0.0001 |
| isoVFA | 0.97418 | -0.22579 | 0.2243 | 0.0095 |

**Diet r2 = 0.68 (P=1e-04)**

#### Charolais (only significant results are presented)

| Pheno | Dim1 | Dim2 | r2 | Pr(>r) |
|---|---|---|---|---|
| DMI_kg_d | -0.68343 | 0.73002 | 0.2414 | 0.0318 |
| total_VFA | -0.31574 | 0.94885 | 0.4083 | 0.0013 |
| C2 | -0.38971 | 0.92094 | **0.4698** | 0.0005 |
| C3 | -0.06482 | 0.99790 | 0.3091 | 0.0098 |
| iC4 | 0.74520 | -0.66685 | 0.3677 | 0.0033 |
| C5 | 0.60220 | 0.79835 | 0.2593 | 0.0284 |
| C6 | 0.04375 | 0.99904 | 0.2516 | 0.0368 |
| C2_C3 | -0.99984 | -0.01789 | 0.3237 | 0.0076 |
| total_protozoa | 0.14976 | -0.98872 | 0.2329 | 0.0357 |
| isoVFA | 0.70708 | -0.70714 | 0.2108 | 0.0510 |
| minorVFA | 0.39788 | 0.91744 | 0.2707 | 0.0277 |

**Diet r2 = 0.21 (P=0.006)**

Pheno: phenotype; LW: live weight; VFA: volatile fatty acid; C2: acetate; C3: propionate, iC4: iso-butryrate; C4: butyrate; iC5: iso-valerate; C5: valerate; C6: caproate; C2_C3: acetate to propionate ratio; iso VFA: sum of iC4 and iC5; minorVFA: sum of iso acids, valerate and caproate.

**Supplementary Table 13. "Animal information Excel file"**

**Supplementary Table 13. Animal information**

**Supplementary Table 14. "Hungate1000 Excel file"**

**Supplementary Table 14.** List of microbial genomes from the Hungate1000 project used in this work

**Supplementary Table 15. Excel file "Supplementary Table 14_Assessment of 324_MAG"**

**Supplementary Table 15.** Evaluation of assembly quality for 324 rumen MAGs

**Supplementary Table 16. Excel file "Supplementary Table 16_rumen-pig-mice_MAG_annotation"**

**Supplementary Table 16.** Taxonomic annotation for MAGs catalog of rumen, pig and mice by CARMA3

**Supplementary Table. Excel file "Supplementary Table 16_DA_KO"**

**Supplementary Table.** Effect of diet on the abundance of KO in the bovine rumen metagenome
